# BioTrendFinder – an interactive web tool for exploring functional drivers in gene- and protein-level bulk omics data

**DOI:** 10.64898/2026.04.12.717932

**Authors:** Alexander G. B. Grønning, Camilla Schéele

## Abstract

The analysis of bulk omics data, such as RNA-seq and proteomics, has enabled numerous biological discoveries. Standard analytical workflows typically comprise dimensionality reduction, group-wise statistical comparisons, functional enrichment analysis, and mapping of molecules to biological networks. Although informative, these steps are often applied independently, limiting integrative interpretation and the efficient identification of functional drivers and candidate targets.

To address these limitations, we developed BioTrendFinder, an interactive web tool for exploring functional drivers in gene- and protein-level bulk omics data. BioTrendFinder employs a sample-ranking strategy to identify significant molecular trendlines that capture expression patterns across ranked sample compositions in dimensionally reduced data. These trends are integrated with statistical results, sample-group metadata and functional information from STRING and eleven bio-ontologies, enabling interactive network-based exploration and the generation of entity-ranked functional modules.

BioTrendFinder’s unique approach and functionalities add additional analytical dimensions to bulk omics data by facilitating the extraction of high-level information from alternative analytical perspectives.

Using previously published proteomics and transcriptomics datasets, we demonstrate that BioTrendFinder supports both hypothesis-driven and exploratory investigations, enabling the prioritization of candidate molecular targets and effectively narrowing the search space for downstream validation steps.

## Introduction

The analysis of bulk omics data, such as RNA-seq and proteomics, has led to numerous pivotal discoveries. Although analytical approaches come in many forms, they typically involve four key steps: (1) dimensionality reduction, using methods such as PCA or UMAP [1], to provide an overview of sample similarities and dissimilarities; (2) group-wise statistical comparisons to identify differentially expressed molecules; (3) enrichment analysis to uncover functional terms associated with significant data segments; and (4) mapping of molecules to biological networks to obtain information about their physical or functional connectivity [2–4].

While each of these steps provides essential insights, they are often treated as separate analytical procedures rather than as interconnected processes capable of revealing higher-order relationships within the data. As a consequence, the overall analytical outcome is frequently limited to descriptions of similarities and differences between conditions, with comparatively limited support for the efficient identification of specific functional drivers and candidate targets. To address these limitations, we developed BioTrendFinder, an interactive web-based tool designed to explore functional drivers in gene- and protein-level bulk omics data.

BioTrendFinder employs a sample-ranking approach to identify sets of significant molecular ‘trendlines’ that capture expression patterns across ranked sample compositions. These expression trends are integrated with contextual information, including sample-group metadata, as well as functional knowledge from STRING [5] and eleven bio-ontologies [6–15]. This integration enables interactive exploration of functional connections through network representations and results in entity-ranked functional modules that highlight molecular contributors that may remain overlooked in conventional analyses.

BioTrendFinder can be used either as a standalone analytical framework or as a complement to standard group-based statistical comparisons and associated analyses. By enabling alternative sample-ranking strategies and downstream analyses, BioTrendFinder introduces additional analytical dimensions for bulk gene and protein data, allowing users to perform question-driven analyses beyond traditional group contrasts. These approaches facilitate the extraction of high-level information from dimensionally reduced data and support interrogation of constructed sample-rankings in relation to underlying expression patterns. The outputs of BioTrendFinder include visual summaries, structured numerical data, and functional annotations, with the computed functional modules serving as a central integrative result of the workflow.

By repurposing previously published proteomics and transcriptomics datasets, we demonstrate how the flexible framework of BioTrendFinder supports both hypothesis-driven and exploratory analyses. By combining expression-based, statistical and annotation data with functional graph information, BioTrendFinder enables the identification of candidate molecular targets of interest and effectively narrows the search space for downstream experimental validation. BioTrendFinder is available at: https://cphbat.shinyapps.io/biotrendfinder/

## BioTrendFinder in general terms

### The main concept of BioTrendFinder

BioTrendFinder was developed to integrate information from dimensionality reduced data (PCA and UMAP), statistical comparisons, sample-group metadata, enrichments analyses and functional connections of molecules. Specifically, the tool was developed to investigate the sample separation along any axis on plots of dimensionality reduced data.

Users can select an axis and thereby generate a ‘*sample-ranking*’, which captures the consecutive arrangement of samples along that axis. For each molecule, a ‘trendline’ is created. Trendlines captures the molecules’ expression patterns across the ranked samples, tracing the molecular expression from the first to the last sample.

Trendlines can be grouped into one or two sets based on increasing or decreasing slope trends. These sets can be statistically filtered, analyzed for sample-group patterns along trendlines and used for enrichment analysis across eleven bio-ontologies.

The molecules from the trendline sets can then be mapped to a functional protein-protein interaction (PPI) network from STRING v. 12, which further can be enriched with the significant terms from the bio-ontologies, enabling interactive exploration of functional connections. The final outputs are functional modules, containing molecules ranked based on expression trends, sample-group metadata and functional connections.

The functional modules highlight the most important expression-based and functional drivers, revealing potential targets of interest, relevant for the specific investigation and the biological context of the analyzed datasets.

In addition, BioTrendFinder also allows users to manually rank samples, create rankings following a molecule’s increasing expression trend, or generate rankings based on predefined sample groups, providing additional flexibility for diverse analyses.

### The flow of BioTrendFinder

BioTrendFinder operates through a sequence of consecutive steps, each represented as a tab within a tabset panel. Most tabs consist of a control panel on the left and a larger main display area on the right.

#### Overview of the tabs

- **Home:** Select one of three analysis tracks (Track 1, Track 2, or Track 3).

- **Upload data:** Upload the required datasets and configure initial parameters.

- **Rank data:** Generate plot-based or manually defined sample-rankings.

- **Analyze:** Compute and evaluate molecular trendlines in the sample-ranked expression data and derive trendline sets.

- **Statistics:** Filter the trendline sets using statistical tests and outlier removal. Visualize significance patterns and sample group conservation.

- **Functional PPI:** Map molecules from the selected trendline sets to STRING’s functional PPI network. The resulting graph can be further enriched using data from the ‘*Enrichment analysis*’ tab.

- **Enrichment analysis:** Perform enrichment analyses and explore the resulting hierarchical functional graphs.

- **Functional module:** Construct functional modules in which molecules are ranked by their inferred importance within the network. Users can also visualize expression data associated with the module’s functional terms.

- **Download:** All generated tables can be downloaded from here.

- **Documentation:** A detailed documentation of all menus, button and checkboxes are provided for every tab-page.

***Note***: all images can be downloaded via PlotLy functionalities or via download buttons at relevant tab-page. The ‘*Download*’ and ‘*Documentation*’ tabs are not shown as part of the BioTrendFinder flow figure (Figure 1), as these serve as supporting elements and not as part of the BioTrendFinder analyses pipeline.

**Figure 1.**
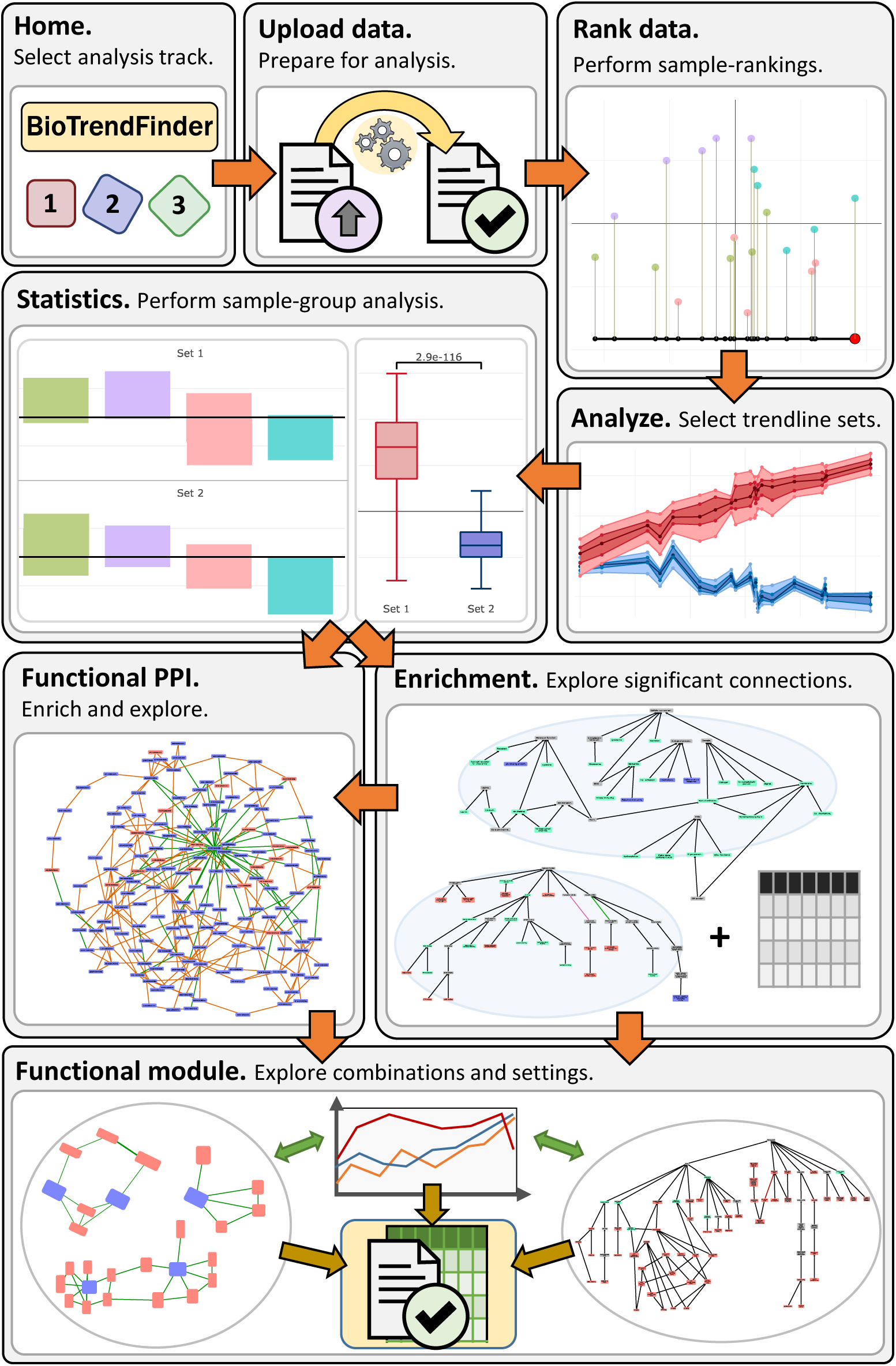
BioTrendFinder flow. A path through all the analytic tabs.

**Figure 2.**
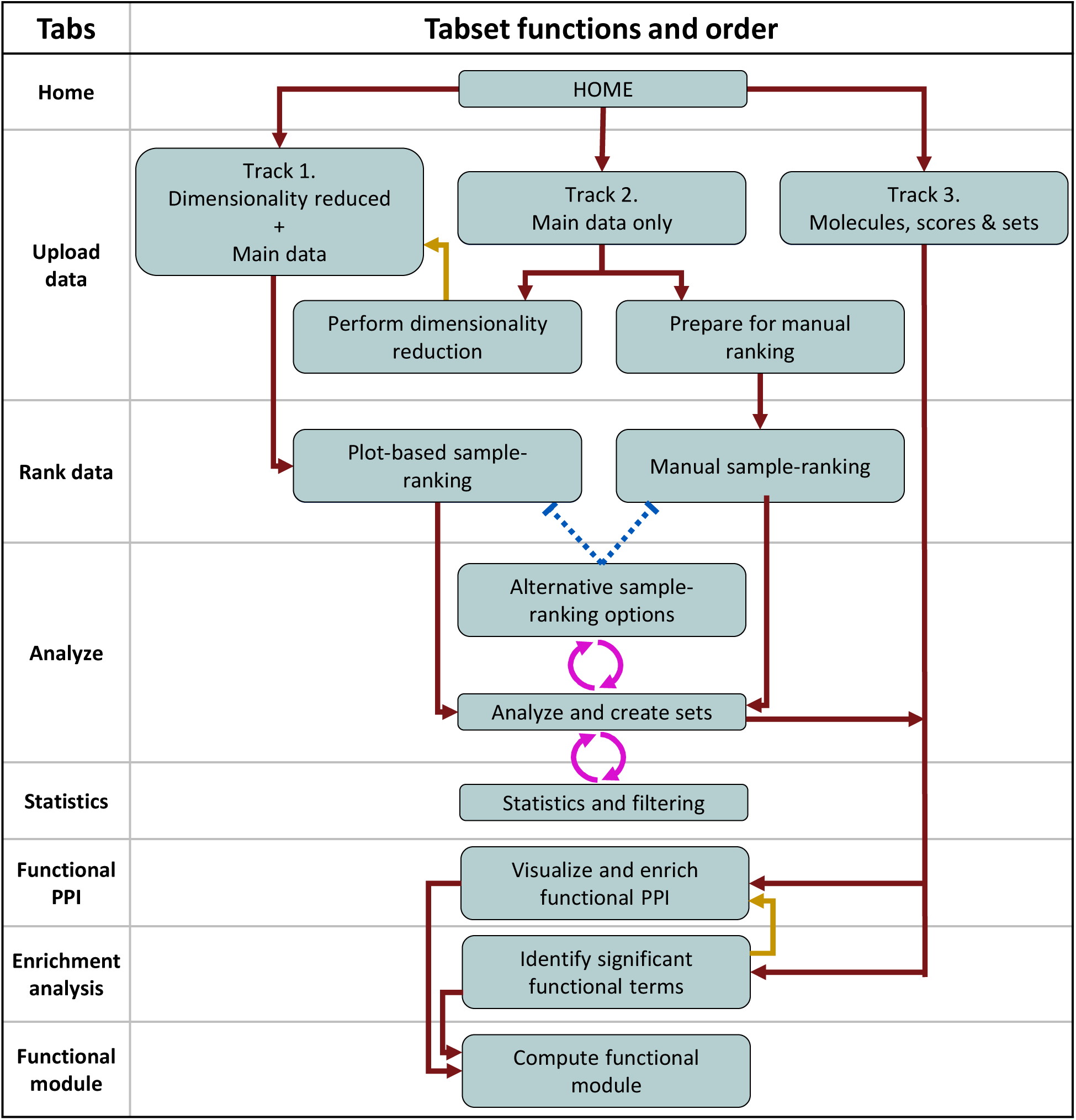
BioTrendFinder flow and tabset connectivity. Options in the ‘Alternative sample-ranking options’ can undo sample-rankings from ‘Plot-based sample-ranking’ and ‘Manual sample-ranking’ indicated with dotted blue lines. Options in the ‘Alternative sample-ranking options’ and ‘Statistics and filtering’ can change the sets defined in ‘Analyze and Create sets’ indicated by pink round arrows. Dark red arrows go down. Dark yellow arrows go up.

### Scope of the manuscript

Comprehensive user documentation is provided within the BioTrendFinder application and is therefore not repeated here. Instead, this manuscript presents an overview of each tab page and its associated functions and options, with the aim of conveying the overall workflow of BioTrendFinder and highlighting its most important functionalities and outputs.

The ‘*Upload data*’ and ‘*Statistics*’ tabs constitute exceptions and are described in greater detail, as proper handling of supported data types and correct application of statistical options are important for effective use of the tool.

In addition, the underlying methods, functions, and algorithms are described in the Methods section. All functions discussed in greater detail there are indicated in the main text by superscript references linking to the corresponding subsections.

### Practical considerations and limitations

While BioTrendFinder provides a flexible and interactive analytical environment, several practical considerations should be noted to ensure appropriate use and interpretation of results.

#### - Extreme values and data normalization

BioTrendFinder assumes that uploaded main datasets have been appropriately normalized to reduce or mitigate the influence of extreme values. Datasets containing uncorrected outliers may lead to suboptimal performance or misleading results.

#### - Limits on trendline set size

The combined number of molecules contained within one or two selected trendline sets is limited to a maximum of 2000 molecules.

#### - Graph visualization constraints

Many functional terms are associated with large numbers of molecules and interactions. To maintain browser performance and usability, the number of edges displayed in the ‘*Functional PPI*’ tab is intentionally limited.

#### - Functional annotations and database coverage

Functional annotations are retrieved from the STRING database (v12), which may not always reflect the most recent updates. As a result, certain functional terms may be outdated. These terms can still contribute to meaningful analyses but are labeled as “missing” when hierarchical graphs are generated in the ‘*Enrichment analysis*’ tab.

Gene Ontology (GO) terms are retrieved in real time via a REST API; consequently, updates to the underlying database may affect result reproducibility over time.

#### - Sample group metadata

Additional analytical insights can be obtained when sample group information is provided and consistently annotated in the input data.

#### - Combining STRING scores with ontology-based connection scores

When combining the STRING scores and the ontology-based connection scores, the two are often not weighted equally, as the STRING scores in BioTrendFinder range from [0.4, 1], whereas the ontology-based connection scores are fractions of the total number of ontology-based connections. As a result, the STRING scores often carry more weight in the combination.

### BioTrendFinder tabs and functionality

The section contains a more detailed description of the BioTrendFinder tabs and associated functions.

#### Home

You can choose between three analysis tracks or choose to reset everything:

#### Track 1 – Dimensionality reduced + main data

You can upload a main dataset and a dataset with dimensionality reduced data. This track is useful if you want to re-analyze data previously generated by BioTrendFinder.

#### Track 2 – Main data only

You can upload a main dataset. At a later step in this track, you can choose to perform your own dimensionality reduction of the main dataset for a plot-based sample-ranking or proceed to the manual sample-ranking option.

#### Track 3 – Upload ID & score data

You can upload a dataset with molecule IDs, associated scores and Set IDs.

#### Reset all

A button allows you to reset BioTrendFinder to start a new analysis from scratch.

### Upload data

You can upload your data and initialize settings for the downstream analyses. The options in the ‘*Upload data*’ tab differ depending on the selected analysis track. BioTrendFinder expects human gene- or protein-level bulk omics data, like RNA-seq, proteomics or peptidomics, as a main dataset. In other words, the molecules in the dataset should relate to a gene or a protein.

BioTrendFinder accepts files with the following suffixes: “.csv”, “.tsv”, “.txt”, “.xlsx” and “.xls”. Once the data are uploaded and the parameters are set, you can proceed to the next tab. The following sections describe the options available under the different analysis tracks.

### Track 1 – Dimensionality reduced + main data

#### Upload of main data

Users can upload a gene-level bulk omics dataset. A selection menu to select the main data column with the most unambiguous molecular identifier is provided. For example, Ensemble gene IDs [16] and UniProt Accession numbers are less ambiguous than gene symbols. Any Ensemble ID version numbers will be removed from the ID list.

#### Upload of dimensionality-reduced data

Users can upload a dataset containing lower-dimensional coordinates for each sample (e.g., PCA or UMAP coordinates). Additional metadata columns, such as sample-group information, may also be included.

Two selection menus are provided:

- **Sample column and coordinates**: Choose the column containing the sample identifiers, and the columns that should be used as the x-axis and y-axis coordinates, respectively.

Three column names in total.

- **Sample-group**s: Select one or more columns containing sample-group information (optional).

#### Track 2 – Main data only

The main data required here are identical to those described for the main data in Track 1. Track 2 provides additional data preparation options:

#### Perform dimensionality reduction

Selecting this option opens a dedicated tab with the following configurable settings:

- **Sample selection:** Choose the columns that define your samples. These will be used as the basis for the dimensionality reduction.

- **Method choice^1^**: Select whether to apply PCA or UMAP.

- **Scaling options:** Specify whether the main data should be zero-centered and scaled to unit variance.

- **Feature filtering:** Select whether to restrict the analysis to the top *X* most variable molecules.

- **STRING filtering:** Optionally limit the dimensionality reduction to molecules that can be mapped to the STRING database.

- **Run reduction:** Click the ‘*Make me’* button to perform the dimensionality reduction. A plot will appear on successful completion.

- **Extract group information^2^:** Click the ‘*Get hover data’* button to automatically derive all possible sample-group annotations from the sample names. These annotations will be used both for plot hover labels and for downstream analyses (***Table 6***).

- **Accept results:** Click the ‘*I’m happy’* button to confirm the dimensionality-reduced dataset and return to the ‘*Upload data’* tab.

#### Prepare for manual selection

Clicking the ‘*To manual ranking*’ button will open a modal window with the following option:

- **Sample selection:** Choose the columns that define your samples. It is possible to select a subset of the total sample list.

***Note***: Remove all columns with numerical values that are not part of your samples, if you wish to generate meaningful group and hover data at a later step.

- **Scaling options:** Specify whether the main data should be zero-centered and scaled to unit variance.

- **Feature filtering:** Select whether to restrict the analysis to the top *X* most variable molecules.

Clicking the ‘*Confirm*’ button will close the modal window and ready you for the next step.

#### Track 3 – Upload ID & score data

Upload a file that contains at least one column with molecule IDs and one column with Set IDs. The ‘*Set ID*’ column may contain up to two distinct characters or symbols, dividing the molecules into a maximum of two sets.

The interface provides three selection menus:

- **Set ID**: Choose the column containing Set IDs.

- **Molecule ID**: Select the most unambiguous molecular identifier.

- **Molecule scores**: Select the score columns associated with each molecule. These scores may be used in later steps

#### Data filtering

When navigating to the next tab (‘*Rank data*’ tab), BioTrendFinder automatically performs the following data filtering steps:

1. Removes version numbers from Ensembl IDs.
2. Maps molecules to STRING IDs and remove entries without a valid STRING ID^3^.
3. Removes rows containing empty (NA) values.
4. Removes rows in which the sample values sum to zero.
5. Removes duplicate molecule IDs (the first occurrence is retained).

### Rank data

BioTrendFinder allows users to generate sample-rankings either manually or based on dimensionality-reduced plots. These rankings are used to generate molecular trendlines for downstream analyses. Once a sample-ranking has been made, click ‘*Proceed!*’ and you will be directed to the ‘*Analysis*’ tab. The following will describe the two different ways to rank your samples.

#### Plot-based sample-ranking

##### Performing rankings

The following describes a small selection of the functions and options located in the main display area:

- **Interactive plot**: Users can click on the plot of the dimensionality reduced data to define an axis along which samples are mapped and ranked consecutively. The current sample-rankings are displayed in a table on the left.

- **Color by sample-groups**: If sample-group data are available, samples can be recolored by group, and group names appear in the legend.

- **Sample deselection**: Individual samples can be deselected using Plotly functions or by clicking legend items.

***Note:*** The plot only captures click coordinates correctly when the page is scrolled to the top.

##### Correlation and connectivity across sample-rankings

The following functions are located in the control panel:

- **Molecule-specific correlation^4^**: Visualize the correlation (Spearman) of a single molecule across a wide selection of possible sample-rankings.

- **Correlation distribution^4^**: Visualize the distribution of Spearman correlations across a wide selection of possible sample-rankings.

- **STRING PPI connectivity distribution^4^**: Visualize the functional connectivity using STRING functional PPI data.

##### Confidence plot and score^5^

When samples are colored according to any of the pre-constructed group annotations, a group confidence plot appears below the clickable plot used to generate the sample ranking. This bar plot visualizes whether the samples within each group are closely clustered along the ranking axis.

#### Manual ranking

Clicking ‘*Manual ranking*’ in the control panel opens a modal window with the following options:

- **Rank samples:** Manually rank your samples according to your desired order.

- **Extract group information:** Click the ‘*Get hover data*’ button to automatically derive all possible sample-group annotations from the sample names. These annotations will be used both for plot hover labels and for statistical analyses (Table 6).

***Note***: This can only be done once.

Clicking ‘*Confirm*’ accepts the manual sample-ranking. After confirmation, the main display area shows a table with the current sample-ranking. From here, you can proceed to the ‘*Analyze tab*’.

### Analyze

In this tab, you analyze and construct your trendline sets. Upon entering ‘*Analyze*’ tab, trendlines for all molecules are automatically generated based on the established sample-ranking.

#### Trendline overview

The main display area shows a plot for individual trendlines and a plot showing the currently selected trendline set – green color indicated an unselected set, red indicates Set 1 and blue indicates Set 2.

A table beneath the plots summarizes characteristics of the selected set. Each trendline is annotated with:

- **Spearman^6^**: The Spearman correlation coefficient of the molecular trendline with respect to the sample-ranking.

- **Slope^7^**: The slope value represents a modified overall increase or decrease of a fitted linear model.

- **Mean_exp**: the mean expression level of the molecule across all samples.

- **Combined_score^8^**: a max-normalized product of the Spearman correlation and the slope.

***Note***: The Combined_score can be redefined to combine any of the available trendline annotation metrics.

#### Defining trendline sets

Three sliders in the control panel and a slider in an ‘*Advanced ranking*’-box (see below) allow the user to define a trendline set by specifying limits for:

- Spearman correlation

- Mean expression

- Slope values

- Combined_score (‘*Advanced ranking*’-box)

Clicking ‘*Apply interval*’ generates a trendline set based on these slider ranges.

Once a satisfactory set has been identified, pressing ‘*Set set*’ in the control panel saves it. Two sets, **Set 1** and **Set 2**, can be defined. At least one set must be created before downstream analyses can be initiated.

#### Advanced ranking

Enabling the ‘*Advanced ranking*’ checkbox in the control panel reveals additional options in the main display area to customize the Combined_score and to set cutoff ranges used for defining a trendline set. Functions for extracting the top or bottom *X* trendlines based on the Combined_score are also provided.

In the ‘*Advanced ranking*’-box, a button labeled ‘*Alternative sample-ranking options*’ opens a modal window that exposes additional ranking strategies (see below).

#### Alternative sample-ranking options

The modal window provides three ranking methods. A molecule must be selected before performing any ranking:

- **Sample values**: Rank any molecule by sample values in increasing order.

- **Group-based**: Rank a selected molecule based on the mean or median of groups of the selected group type in increasing order.

- **Manual**: Manually rank a selected molecule according to groups of the selected group type in a user-defined order.

***Note*:** For the **group-based** and **manual** options, users can choose to collapse groups into a single data point. This preserves the group order, but allows sample-rankings within the groups to differ across trendlines, making the ranking of each trendline individual (similar to method the **group-based** option) rather than identical across all trendlines.

### Statistics

This tab provides tools to identify significant trendlines and remove insignificant ones, strengthening the defined trendline sets and ensuring that trendlines exhibit differentially expressed segments. The underlying assumption is that each trendline represents either an increasing or decreasing trajectory. BioTrendFinder therefore assesses differences between end-segments to detect significant contrasts. The statistical filtering can be accepted to remove insignificant trendlines from the sets. In addition, the trendline sets (Set 1 and Set 2) themselves can be statistically compared, if the two trendline sets have been made.

***Note***: Accepting filtering may change the number of trendlines in a set. Statistical filtering should therefore be finalized before downstream analyses if deemed relevant.

Comparing two entire trendline sets is not always meaningful, as sets with highly distinct expression trends may nonetheless be drawn from the same underlying distribution.

### Modes of comparison

The control panel provides an interactive menu controlling whether segments or groups are used for the statistical comparisons.

#### Section-based comparisons^9^

Trendlines are divided into a user-defined number of smaller segments. BioTrendFinder compares all combinations of left-side vs. right-side end-segments using the Welch test. In addition, for each comparison the log2 fold change (log2FC) and a pi-score are computed.

#### Group-based comparisons^9^

If ‘alternative sample-ranking options’ (any group-based ranking type) have been applied, users may compare combinations of left and right group-segments using the same statistical framework (Welch test, log2FC, and pi-scores).

#### The statistics table

BioTrendFinder automatically identifies the most significant comparisons and performs a FDR correction. Results are shown in a statistics table in the main display area, where new columns are introduced:

- **pval:** the lowest p-value given the settings of **percent included** and **left/right balance**

(see ‘General and advanced statistics’ below).

- **fdr:** FDR correction of the p-values using all items displayed in the statistics table.

- **log2FC:** Log2 fold change of the compared trendline segments.

- **pi_score:** the pi-score (pi-value [17]) of the compared trendline segments.

- **left_right**: the number of the left-side and right-side data points included in the comparison.

- **percent_included**: the percentage of samples participating in the comparison (1 indicates full inclusion).

***Note***: Because all p-values in the statistics table are included in the FDR correction, the resulting FDR values for items in, e.g., Set 1 may differ when this set is selected alone compared with when both sets are selected.

#### Statistics for individual molecules or the entire set

Users may examine the statistics in detail at either an ‘*individual molecule*’ level or an ‘*entire set*’-level. The display can be switched by clicking relevant checkboxes located in the control panel.

#### Individual molecule

Select a molecule from the statistics table and click ‘*Molecule significance*’ to view all comparisons made in an interactive plot. Two plots appear beneath the table in the main display area:

- **Significance of comparisons**: all comparisons as selectable data points.

- The y-axis of the left plot is adjustable through a selection menu in the control panel. The default y-axis value is −𝑙𝑜𝑔10(𝑝𝑣𝑎𝑙).

- **Trendline and boxplot**: the corresponding compared sections of a trendline and an accompanying boxplot showing segment differences.

A button labeled ‘*Set this left/right balance for all’* right below the statistics table applies the chosen comparison structure to all molecules in the selected set.

#### Entire set

Clicking ‘*Entire set significance*’ button visualizes the distribution(s) of the selected set(s). Only the trendlines that passes the given alpha and log2fc threshold are included on the visualizations that are displayed below the statistics table:

- **Sample-ranked expression distribution**: Distribution of expression values across the sample-ranking.

- **Overlap and boxplot**: An overlap plot showing the segments detected in the significant trendlines, accompanied by a boxplot comparing left- vs. right-side segments.

If both sets are selected, the boxplot will illustrate the distribution of Set 1 vs. Set 2.

When the ‘entire-set’ checkbox is selected, an option called ‘bundle-up’^10^ becomes available. Clicking this option changes how the trendline distributions are displayed, emphasizing the overall patterns (the trends) within each set of trendlines.

If sample-group information is available, an additional selection menu appears, enabling group-level visualization:

- **Group distribution**: distribution of group membership along the x-axis.

- **Group inclusion summary**: percentage of each group included in left and right segments.

***Note***: the group data can be used to evaluate the groups division along the trendlines^11^.

#### General and advanced statistics

##### Set selection

A selection menu allows users to display the statistics for ‘*Set 1*’, ‘*Set 2*’ or ‘*Both*’. Changing the menu updates the statistics table.

##### Statistical thresholds

Controls for ‘*alpha*’ and ‘*log2FC*’ thresholds are provided. Increasing alpha and lowering the log2FC threshold relaxes significance requirements (setting alpha = 1 and log2FC = 0 includes all trendlines).

##### Advanced statistics and outlier removal^12^

The control panel contains an ‘*Advanced statistics*’ checkbox. Enabling this option opens a menu for outlier filtering where users can specify:

- **Max number of items to remove:** The maximum number of items to remove from trendline segments when compared.

- **Min/max ratio:** The maximum allowed ratio between the highest and lowest values across the compared trendline sections. A value of 1 ensures that, e.g., for an increasing trendline, the highest value in the left-side segment does not exceed the lowest value in the right-side segment.

A red button will initiate the removal function, and Clicking ‘*Apply*’ updates the statistics table.

##### Percentage included and left/right balance

Clicking the ‘*Show/hide inclusion box*’ link located in the control panel will reveal a menu where the sizes of the compared trendline segments can be controlled. The menu contains two fields for numerical inputs and a button to accept any changes. Both numeric input fields accepts values in the range [0,100]:

- **Min % of samples included:** this value controls the number of samples included in the comparisons. A value of 100 ensures that all samples are used for the comparisons.

- **left-right balance (%):** this value refer to the one obtained by taking size of the smallest end segment in a trendline and dividing by the size of the largest end segment. A value of 100 ensures that the compared segments are perfectly balanced (have identical sizes).

Clicking the ‘*Update data’* button will update the statistics table.

##### Accept statistical filtering

Two checkboxes at the bottom of the control panel allow users to confirm statistical filtering for Set 1 and Set 2. Once checked, statistics for that set become locked and cannot be recomputed. ***Note***: Statistical filtering can be applied to a set only after entire set statistics have been computed.

### Functional PPI

This tab provides tools for exploring molecules from the defined trendline sets within an interactive functional PPI network environment.

#### STRING functional PPI visualization and general functions

Clicking the “Visualize STRING network” button maps the molecules to the STRING v12.0 database and updates the interactive VisNetwork-based graph. Edges can be clicked to display the underlying interaction score. Users may also modify and customize the minimum required STRING interaction score according to the STRING scoring scheme [18].

Additional network control buttons allow users to:

- **Set focus**: View the individual trendline sets separately.

- **Toggle singletons**: Toggle the presence of unconnected nodes.

- **Physics control**: Toggle network physics for improved exploration.

***Note***: only when physics has been disabled will edges appear.

- **Label change**: Adjust node labels based on alternative molecule ID columns from the uploaded main dataset.

- **Louvain communities**^13^: users can perform Louvain community detection.

- **Node selection:** Users allowed to select Louvain communities (if computed) and/or any subset of nodes for subsequent functional graph enrichment or functional module construction.

#### Enriching functional graph^14^

If enrichment analyses have been completed in the ‘*Enrichment analysis’* tab, an additional menu becomes available. This menu enables users to select functional terms across all supported bio-ontologies, which are then used to enrich and extend the functional graph. In doing so, biological annotations are integrated directly into the network structure. For each functional term, BioTrendFinder adds edges between all nodes associated with that term^15^.

##### Advanced annotation selection

Button called ‘*Advanced selection’* opens a menu where all available significant terms can be manually selected. Here, users can filter the functional terms by different parameters that relate to the naming or underlying statistics. A parameters called ‘*Set entropy*’^16^ has been included when two trendline sets are involved in the analysis.

***Note***: this menu is also available in the ‘*Functional module’* tab, once significant functional terms has been found.

### Enrichment analysis

This tab provides tools for performing enrichment analyses across multiple biological annotation resources. All supported ontologies are searched automatically, and when multiple trendline sets are defined, the associated molecules are treated as a single combined set. This enables BioTrendFinder to identify functional terms shared among the selected molecules. Only terms that pass the false discovery rate (FDR) cutoff are displayed (default value: 0.01)^17^.

#### Bio-ontologies^18^

The following bio-ontologies and databases are queried:

1. Gene Ontology – Biological Process [6, 7]
2. Gene Ontology – Cellular Component [6, 7]
3. Gene Ontology – Molecular Function [6, 7]
4. Annotated Keywords (UniProt) [8]
5. Disease gene associations (DISEASES) [9]
6. Human Phenotype Ontology (Monarch Initiative) [10]
7. KEGG pathways [11]
8. Reactome pathways [12]
9. WikiPathways [13]
10. Subcellular localization (COMPARTMENTS) [14]
11. Tissue expression (TISSUE) [15]

#### Ontology graph visualization^19^

Upon successful computation, users can visualize bio-ontology–specific significant terms as hierarchical graphs. The graph is returned as a connected component; additional insignificant nodes required for maintaining graph connectivity are displayed as grey nodes. Significant terms are color-coded: red for Set 1, blue for Set 2, and green for terms associated with both sets.

Clicking any node updates the enrichment table at the bottom of the page.

***Note*:** Some annotation terms originate from STRING v12.0 and may be obsolete or missing in current ontology definitions. Such terms are excluded from the graph but are still listed in the enrichment table. A dedicated menu allows users to display both added (grey) nodes and missing terms.

#### Functional module

This tab becomes available once the molecules have been mapped onto the STRING functional graph and/or an enrichment analysis has been completed.

BioTrendFinder constructs an integrated functional graph using either the STRING-derived edges and/or ontology-based term connections. From this graph, molecule-level scores are computed, enabling the identification of functional modules and the ranking of their molecular constituents.

***Note***: it is possible to add unconnected molecules to the functional module as well. These will have zero values in all score metrics relating to the functional graph (degree_centrality, edge_weight_fraction and node_importance).

#### Functional module table

The resulting table introduces several new columns:

- **Nodes**: identifiers of the molecules included in the module.
- **Degree_centrality^20^**: normalized estimated degree, adjusted for the number of components within the module.
- **Edge_weight_fraction^21^**: the fraction of the total edge weight connected to each node.
- **Node_importance^22^**: a composite score summarizing node impact based on connectivity and total edge weight contribution.
- **FM_score^23^**: Functional module score, the average of all selected score types used in the module computation.
- **Terms**: identifiers of the functional terms associated with the node set.

The **FM_score** can be recalculated to incorporate trendline-derived metrics and expression data, statistical results and annotation-based scores from upstream steps. This score determines the ranking of molecules within the module: higher values indicate stronger functional prominence within the context of the selected sample-ranking and chosen functional terms. The combined use of expression dynamics, statistical significance, and graph structure enables the identification of molecules that may represent novel targets of interest.

#### Visualizing functional terms in module

The functional module table can be swapped with a module-specific enrichment table listing all terms represented within the module. Selecting a term highlights the expression patterns of the molecules associated with it, enabling direct inspection of functional coherence and sample-dependent behavior.

## Analysis examples

All analysis examples presented here are framed in the context of adipose tissue and metabolic phenotypes to highlight BioTrendFinder’s ability to identify biologically relevant targets.

### Exploratory analysis of proteomics secretome

To assess the utility of BioTrendFinder, we analyzed a previously published proteomics dataset [19]. The dataset comprises mass spectrometry analysis of conditioned media from adipose progenitor cells that were isolated from human brown adipose tissue (BAT) or white adipose tissue (WAT). The cells were differentiated in vitro and were either untreated (woNE) or treated with norepinephrine (NE). The NE treatment aimed to simulate sympathetic activation of adipocytes upon cold exposure. Data and results can be downloaded from https://github.com/scheelelab/BioTrendFinder_results_and_data.

We performed an exploratory analysis of the dataset to address the following questions:

1. Which proteins in the cell media contribute to the separation of samples along the first principal component of the PCA?
2. Do any of these proteins exhibit consistently increasing or decreasing expression trends?
3. Are the resulting proteins associated with common functional terms, and can this information be used to identify novel proteins relevant to the dataset and associated conditions?

#### Steps with BioTrendFinder

These are the steps taken in the different tabs:

1. **Home**: Selected the “Main dataset only” analysis track (Track 2).
2. **Upload data**: Uploaded the data, and selected the column with UniProt accession IDs, as the column with primary molecular identifiers. We then computed a new PCA plot and associated data, and generated group data based on the sample names of the dataset.
3. **Rank data**: After the automatic dataset filtering, the dataset had shrunk from 1065 to 1017 rows (proteins). We then generated a ranking-line that followed the x-axis of the PCA plot (PC1, **Figure 3**A).
4. **Analyze**: For Set 1, trendlines were selected based on a Spearman correlation coefficient

**Figure 3.**
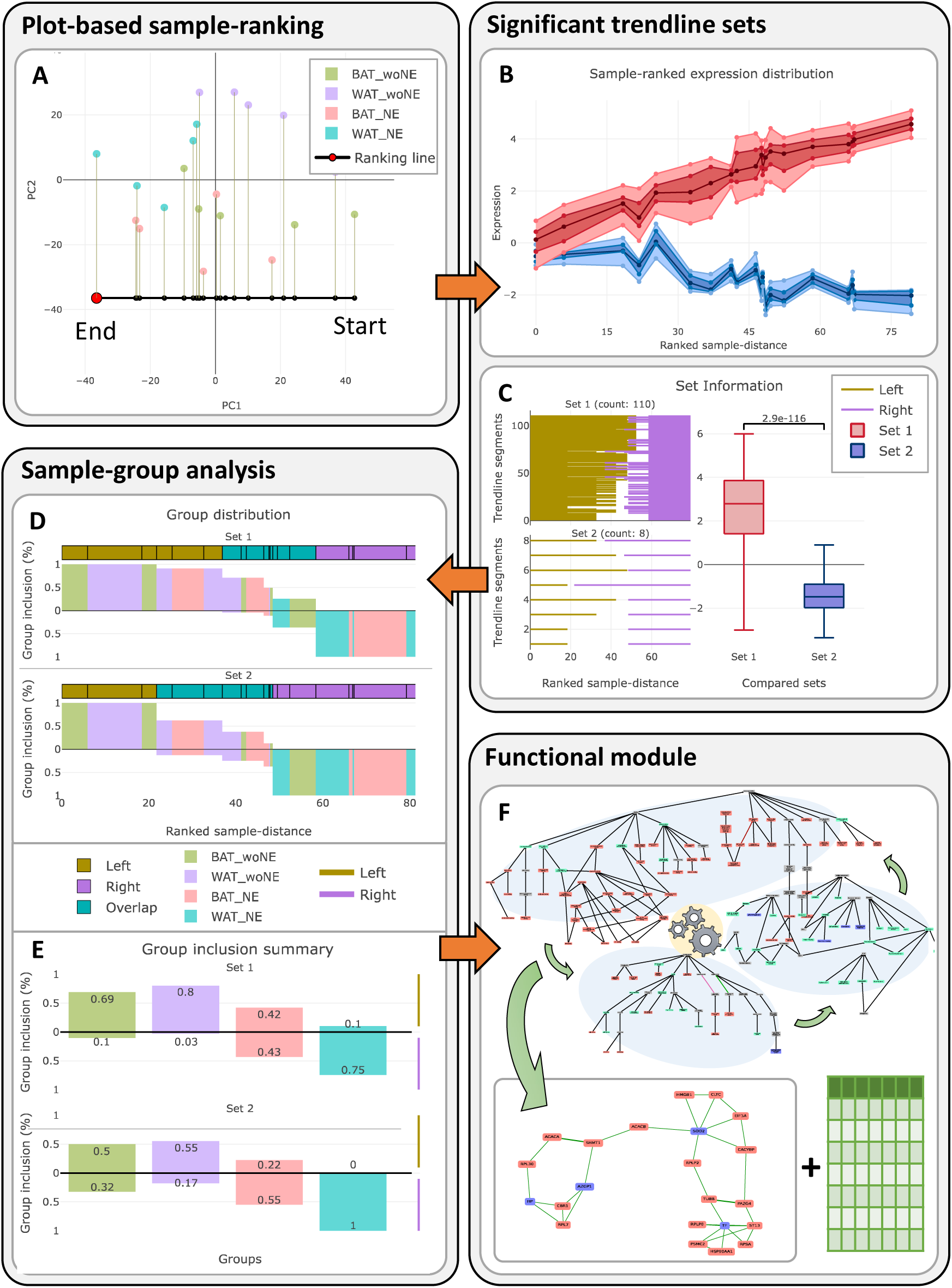
**BioTrendFinder workflow for exploratory analysis**. The process begins with plot-based sample ranking, in which a ranking line is derived from a PCA plot (A). Significant trendline sets are then defined (B), along with an associated statistical overview (C). A sample-group analysis reveals group distributions along the trendline sets (D) and provides a summary of sample-group divisions (E). Finally, a functional module is computed using bio-ontologies (F).

≥ mean + standard deviation of the correlation distribution and a slope value ≥ mean + standard deviation of the slope distribution. For Set 2, trendlines were selected using thresholds of Spearman correlation ≤ −0.6 and slope ≤ −1.The sizes if Set 1 and Set 2 were 114 and 8, respectively.

1. **Statistics**: Identified significant trendlines for each set individually. We divided the trendlines into 10 segements (default), used an FDR alpha=0.01 and a log2fc threshold of

0.172 (computed automatically per default). We checked the box ‘log2 transformed already’. For the visualization of the significant trendline sets, we checked the ‘Bundle up’ checkbox. The final sizes if Set 1 and Set 2 were 110 and 8, respectively (**Figure 3B, C**). For the final trendline sets, we conducted a sample-group analysis detecting sample-group division patterns (**Figure 3D, E**).

1. **Enrichment analysis**: We performed enrichment analysis across all available bio-ontologies with an FDR correction filter at 0.01.
2. **Functional module**: we identified functional terms where proteins from both Set 1 and Set 2 closer to equally involved, by setting the ‘*set entropy*’ parameter to 0.5 in the ‘advanced settings’ menu. This resulted in 29 significant functional terms across GO-terms (BP, CC), Reactome, COMPARTMENTS and TISSUES, which connected 88 proteins (80 from Set 1 and 8 from Set 2). The module was ranked using: degree_centrality, spearman, pi_score, and percent_included. All proteins were included in the module (**Figure 3F**).

We downloaded all images at the different relevant tab-pages and downloaded all the needed data at the ‘*Download*’ tab.

#### Group division scores

The left side of the trendlines in Set 1 (low-expression region) exhibited moderate and strong group-division trends for **BAT_woNE** (score = 0.58) and **WAT_woNE** (score = 0.72), respectively, relative to the null model (Table 8). In contrast, the right side of the same trendlines (high-expression region) showed a moderate/strong division trend for **WAT_NE** (score = −0.64). **BAT_NE** samples were evenly distributed between the left and right sides of the trendlines, indicating no preferential association with either the low- or high-expression regions.

In **Set 2**, most group division trends were negligible to weak (**BAT_woNE**: 0.175; **WAT_woNE**: 0.375; **BAT_NE**: −0.325). An exception was **WAT_NE**, which showed a division score of −1 on the right side, indicating complete inclusion of **WAT_NE** samples in the low-expression region of the trendlines.

#### Interpretation

When following the sample-ranking along PC1, the proteins exhibiting significantly increasing trendlines (Set 1), showed clear sample-group division patterns. Specifically, **BAT_woNE** and **WAT_woNE** samples were generally associated with lower expression levels, whereas **WAT_NE** samples tended to occupy higher expression regions.

Proteins with significant decreasing expression trends (Set 2) were fewer and displayed more ambiguous group-division patterns. Nevertheless, these proteins consistently exhibited lower expression levels in **WAT_NE** samples compared to the other samples.

#### Functional module – result and discussion

The resulting module integrates functional graph information from bio-ontologies, protein expression trends and trendline-derived statistics (degree_centrality, spearman, pi_score, and percent_included). The top five proteins from Set 1 and Set 2 can be seen in **Table 1** and **Table 2**, respectively.

**Table 1.** Top five molecules of the increasing trendline set (Set 1).

| Rank | nodes | Gene name | Reported associations with adipose tissue and metabolic phenotypes | Ref. |
| --- | --- | --- | --- | --- |
| 1 | P50990 | CCT8 | Delaying aging in *C. elegans*, upregulated during brown adipose tissue thermogenic lipolysis, higher number of small extracellular vesicles in T2D. | [4, 29, 30] |
| 2 | P39023 | RPL3 | Higher expression in ‘low heat loss mouse model’ vs. ‘high heat loss mouse line’ in BAT and hypothalamus. | [31] |
| 3 | P07900 | HSP90AA1 | Involved in browning. General modulation of wound healing, cell motility, and inflammation. | [27, 28] |
| 4 | P43490 | NAMPT | Insulin sensitivity, impaired by aging. Obesity-decreased adipose tissue expression, diet-induced weight loss increases adipose tissue expression. | [25] |
| 5 | P35579 | MYH9 | Obesity and adipogenesis regulation. | [32, 33] |

**Table 2.** Top five molecules of the decreasing trendline set (Set 2).

| Rank | nodes | Gene name | Reported associations with adipose tissue and metabolic phenotypes | Ref. |
| --- | --- | --- | --- | --- |
| 8 | P25311 | AZGP1 | Lipid metabolism. Decreased expression in adipose tissue and liver and lower plasma protein levels in obese mice. | [21, 34] |
| 16 | P00738 | HP | Inflammatory and adiposity marker. Higher expression in WAT associated with obesity, increased blood concentration in obese humans. | [24, 35, 36] |
| 45 | P04179 | SOD2 | Adipogenesis, immunosuppression, diabetic wound healing. | [37, 38] |
| 51 | P02787 | TF | Insulin resistance, type 2 diabetes. | [22, 39] |
| 52 | B3KW70 | MFAP5 | Adipose tissue inflammation, adipogenesis. | [23, 40, 41] |

The top five proteins in the decreasing set (Set 2) (enriched in the untreated condition) include several known adipokines. These proteins are associated with the extracellular secretome involved in adipocyte homeostasis, extracellular matrix organization, and systemic metabolic buffering, indicating that the decreasing set shows high abundance of proteins linked to a basal extracellular maintenance program. In addition, four out of the five proteins have been directly implicated in metabolic pathologies: AZGP1 has been reported to counteract the effects of obesity [20, 21], and TF has been associated with a protective role against the development of insulin resistance [22]. In contrast, MFAP5 is linked to inflammation and extracellular matrix remodeling [23], while HP deficiency has been reported to confer protection against obesity-associated comorbidities [24].

The top five proteins in the increasing trendline set contain intracellular proteins not directly associated with secretion. These include CCT8, a cytosolic chaperonin component; RPL3, a ribosomal protein; and MYH9, a cytoskeletal motor protein.

NAMPT on the other hand is known to act both intracellularly and as a secreted adipokine, and has been found to be central in many adipose-related processes including various aging-related and metabolic disorders [25, 26]. HSP90AA1 (Hsp90α) is abundantly expressed can be secreted extracellularly in response to a variety of cellular stresses to facilitate adjustment to the stress [27]. Interestingly, a recent study has implicated it in browning upon secretion from skeletal muscles during training [28], which calls for a similar study in an adipose setting.

Altogether, looking at the top five proteins of the increasing set seems to reflect a stress- and remodeling-related module. The distinct patterns of the increasing and decreasing modules suggest a transition from a classical adipocyte secretory profile toward an alternative extracellular signature enriched in intracellular proteins. Whether this signature reflects cellular stress induced when moving from BAT/WAT woNE towards WAT treated with NE, adaptive signaling, or a combination of both remains to be determined. Regardless, by using BioTrendFinder we identified a pattern which went unnoticed in the initial analysis, and which should be considered in future studies.

### Extracting obesity ameliorating factors

In this study we used a previously published bulk RNA-seq dataset based on subcutaneous abdominal adipose tissue biopsies from 20, 20 and 15 human adults categorized as metabolically unhealthy obesity (MUO), metabolically healthy obesity (MHO) and metabolically healthy and lean (MHL), respectively. Data and results can be downloaded from https://github.com/scheelelab/BioTrendFinder_results_and_data.

For this test case we had a more precise question to direct at BioTrendFinder:

1. Can increasing and decreasing trendlines from MUO to MHO to MHL samples be established?
2. Are any of the resulting genes annotated as secreted and can they be used to ameliorate some of the unhealthy effects of the MUO phenotype?

#### Steps with BioTrendFinder

1. **Home**: Selected the “Main dataset only” analysis track (Track 2).
2. **Upload data**: After dataset uploaded, we selected the column with Ensembl gene IDs as the column with primary molecular identifiers. We proceeded the manual ranking option.
3. **Rank data**: After the automatic dataset filtering, the dataset had shrunk from 34259 to 19240 rows (transcripts). All samples were selected at random and sample-group information was extracted from the column names.
4. **Analyze**: The ‘Alternative sample-ranking option’ was selected. In this menu we chose to ‘collapse groups’ and the group means were used as representing values. The sample-groups were selected in the following order: MUO, MHO, MHL (**Figure 4**). For Set 1, we extracted the trendlines that had a combined_score ≥ mean + 2*standard deviation of the combined_score distribution and Spearman correlation score == 1. For Set 2, we extracted the trendlines that had a combined_score ≤ mean - 2*standard deviation of the combined_score distribution and Spearman correlation score == -1. The size of Set 1 was 197 and the size of Set 2 was 684.
5. **Statistics**: The statistical tests were based on the group-division scheme, and the settings were modified to include 100% of the samples. The significant trendlines for the sets were defined simultaneously using the default FDR (0.01) and log2fc (0.046) thresholds. The resulting set sizes were 170 and 644 for Set 1 and Set 2, respectively. For the final trendline sets, we conducted a sample-group analysis detecting group division patterns (**Figure 3**B).
6. **Functional PPI**: The genes were mapped to the functional PPI from STRING v. 12.0 to allow for the usage of STRING’s functional connections in the ‘Functional module’ tab.
7. **Enrichment**. Significant functional terms were identified using the default settings.
8. **Functional module**: The functional connections from STRING were utilized together with the term “Secreted” (KW-0964 from Uniprot Keywords) in the advanced settings, which filtered away all molecules not associated with the annotation. The resulting functional module contained 166 genes. The functional module score (FM_score) was defined using the parameters node_importance, combined_score, fdr, log2fc and percent_included (**Figure 3**C).

#### Group division scores

The left side of the trendlines in **Set 1** (low-expression region) exhibited very strong and strong group-division trends for **MOU** (score = 1) and **MHO** (score = 0.72), respectively, relative to the null model (Table 8). The right side of the same trendlines (high-expression region) showed a very strong division trend for **MHL** (score = −1).

**Figure 4.**
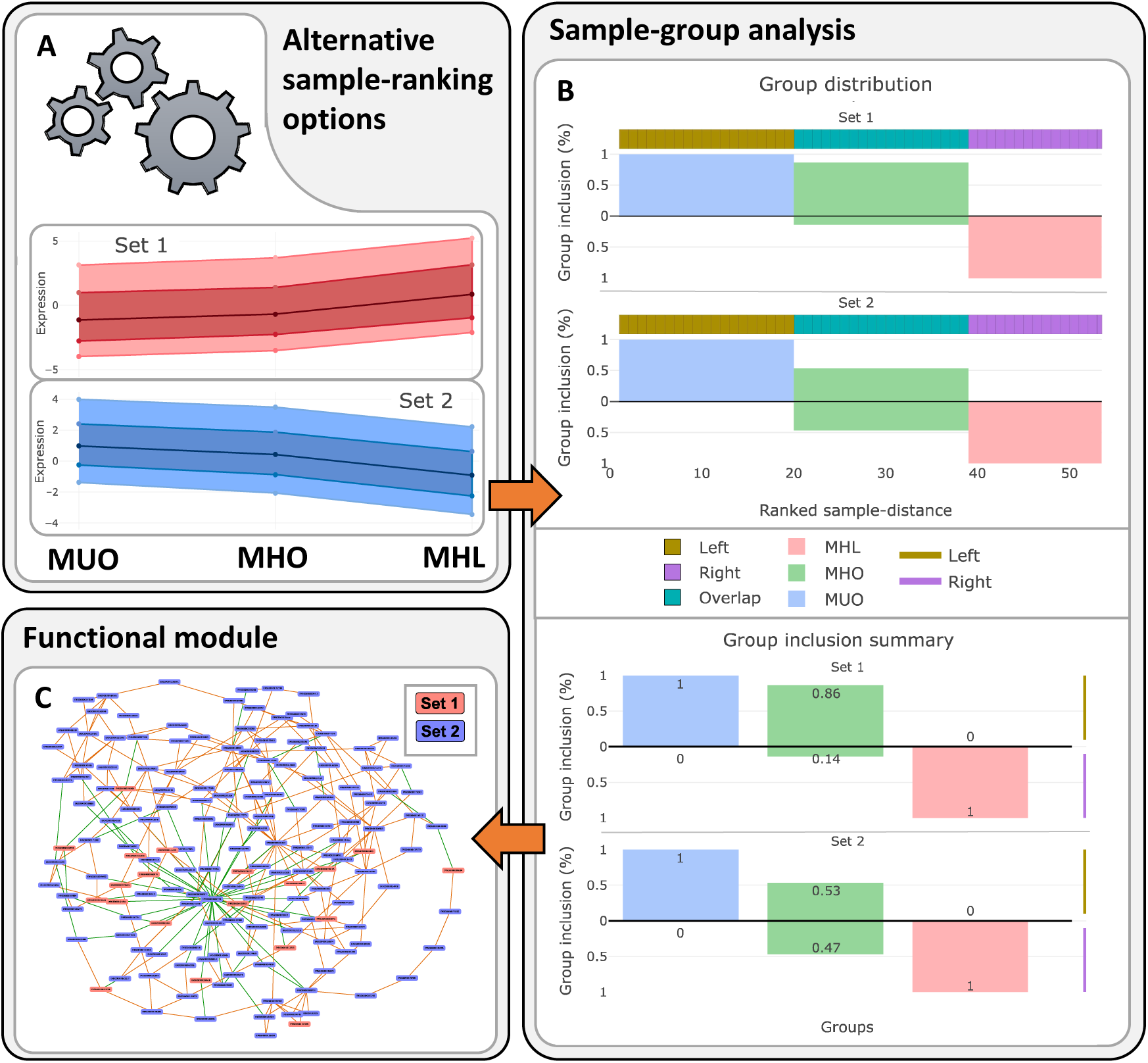
**BioTrendFinder flow summary of metabolic analysis**. The process begins with alternative sample ranking, which collapses group samples into single representative points (A). Next, a sample-group analysis is performed (B), followed by computation of a functional module (C).

In **Set 2**, the left side of the trendlines in **Set 2** (high-expression region) exhibited very strong group-division trends for **MOU** (score = 1). The right side of these trendlines (low-expression region) showed a very strong division trend for **MHL** (score = −1). Here, **MHO** showed none to very weak division trends, indicating no preferential association with either the low- or high-expression regions.

#### Interpretation

In the sample-ranking going from the MUO to MHO and ending at the MHL samples, the transcripts in the increasing trendline set (Set 1) exhibited clear sample-group division patterns, with MUO and MHO samples predominantly occupying regions of lower expression, while MHL samples were enriched in regions of higher expression.

For the decreasing trendline set (Set 2), the MUO samples trended to occupy the high-expression region and the MHL samples were largely located in the low-expression area. The MHO samples were approximately evenly distributed between the high- and low-expression regions.

#### Functional module – result and discussion

The resulting module integrates functional graph information from STRING, protein expression trends and trendline-derived statistics (degree_centrality, spearman, pi_score, and percent_included) (**Table 3**, **Table 4**). Among the top five transcripts in both the increasing and decreasing sets are candidates with potential therapeutic relevance in the context of obesity and associated metabolic disorders.

**Table 3.** Top five molecules of the increasing trendline set (Set 1).

| Rank | nodes | Gene name | Reported associations with adipose tissue and metabolic phenotypes | Ref. |
| --- | --- | --- | --- | --- |
| 1 | ENSG00000152785 | BMP3 | Associated with obesity and being predictive of obesity development in mouse models. | [45, 46] |
| 8 | ENSG00000145321 | GC | Associated with obesity; genetic ablation in mice improved metabolic health. | [47, 48] |
| 14 | ENSG00000134548 | SPX | Ameliorates obesity-induced metabolic disorders by improving WAT browning. Stimulates lipogenesis and inhibits adipogenesis, lipogenesis and glucose uptake in human and murine 3T3-L1 adipocytes. | [49, 50] |
| 32 | ENSG00000206384 | COL6A6 | Vaccine alleviates atherosclerosis via regulation of the immune system and lipid metabolism. | [43] |
| 36 | ENSG00000160862 | AZGP1 | Involved in lipid metabolism; reduced expression in adipose tissue and plasma in obese mice. | [21, 34] |

**Table 4.** Top five molecules of the decreasing trendline set (Set 2).

| Rank | nodes | Gene name | Reported associations with adipose tissue and metabolic phenotypes | Ref. |
| --- | --- | --- | --- | --- |
| 2 | ENSG00000060718 | COL11A1 | Upregulated in obesity-induced adipose tissue expansion. Proposed PC and NAFLD biomarker. | [51, 52] |
| 3 | ENSG00000069482 | GAL | Linked to increased fat ingestion and obesity exacerbation. Potential therapeutic target | [53, 54] |
| 4 | ENSG00000133063 | CHIT1 | Higher expression by adipose tissue macrophages in overweight subjects and T2D patients. Inhibition attenuates metabolic dysregulation | [44, 55] |
| 5 | ENSG00000198759 | EGFL6 | Increased expression in obesity. Involved in obesity-related AT dysfunction in children | [56, 57] |
| 6 | ENSG00000134028 | ADAMDEC1 | Intestinal mucosa inflammation and upregulation in obese rats. Lower expression associated with Crohn disease. | [58] |

Notably, SPX from the increasing set (Set 1) has been reported to suppress appetite, whereas GAL from the decreasing set (Set 2) is associated with increased fat intake. Accordingly, SPX agonists and GAL antagonists may represent promising therapeutic strategies [42].

AZGP1 from Set 1 has been reported to increase leptin sensitivity and may therefore represent a potential therapeutic target for obesity-associated metabolic disorders [20]. In addition, a COL6A6-based vaccine has been reported to alleviate atherosclerosis [43].

In contrast, A CHIT1 inhibitory drug OATD-01 has reached phase II clinical trials in patients with lung sarcoidosis and has shown promise as a therapeutic agent for metabolic dysfunction-associated steatohepatitis (MASH) [44]. For the remaining transcripts in the increasing and decreasing sets, an important avenue for future research is to determine whether any may also serve as potential therapeutic targets.

When relating the aforementioned genes to the transcriptomic data, the observed effects and proposed therapeutic implications align with the predefined sample groups. Specifically, transcripts with higher expression in MHL compared to MUO samples were enriched for factors associated with metabolic health and obesity amelioration, such as SPX and AZGP1. Similarly, transcripts more highly expressed in MUO relative to MHL samples were associated with factors linked to obesity and related comorbidities.

## Discussion

In this work, we present a focused walk-through of the functionalities and analytical options available in BioTrendFinder, an interactive web tool for exploring functional drivers in gene- and protein-level bulk omics data. The presentation is intentionally weighted toward explaining the rationale, core functionalities and outputs of the different tab pages, as well as elaborating on the underlying methodology and algorithms. A comprehensive description of individual interface elements, including buttons, checkboxes and configuration options, is provided in the ‘*Documentation*’ tab on the BioTrendFinder website.

BioTrendFinder can be used either as a standalone analytical framework or as a complement to standard group-based statistical comparisons and associated analyses. By enabling novel sample-ranking strategies and downstream analyses, BioTrendFinder introduces additional analytical dimensions for bulk gene- and protein-level data, allowing users to pose alternative, question-driven analyses beyond conventional group contrasts. Through these approaches, users can extract high-level information from dimensionally reduced data or interrogate designed sample rankings in relation to underlying expression patterns. The outputs of BioTrendFinder range from visual summaries to structured numerical data and functional annotations, with the computed functional modules representing a central integrative result of the workflow.

All molecules are mapped to STRING IDs (v12), and unmappable molecules are removed, resulting in some data loss. Despite this, BioTrendFinder is able to uncover novel, biologically meaningful patterns, highlighting its robustness in identifying previously unrecognized molecular associations.

Two test cases based on bulk proteomics and RNA-seq data were included to demonstrate the utility of BioTrendFinder as a robust hypothesis generator and tool for target exploration and identification. For all identified functional modules, we only inspected a subset of the molecules identified in the trendline sets, leaving enormous room for further analysis and exploration.

In the first test case, we highlight the trend-finding capabilities of BioTrendFinder and its ability to extract high-level information, such as sample-group trends, from a PCA-based representation of the data. In the second test case, we demonstrate how the inherent features of the tool can be used to generate user-defined sample-rankings and to identify molecules that may address the specific analytical questions posed at the outset. Across both analyses, we identified sets of highly functionally connected and inversely regulated molecules with established or emerging roles in adipose tissue physiology, metabolism, and related disorders, consistent with the biological context of the analyzed datasets.

Interestingly, comparison of the functional modules computed from the two test cases revealed an overlap, as both AZGP1 and MFAP5 were identified in each analysis. In the proteomics study, AZGP1 and MFAP5 were expressed at lower levels in WAT_NE samples compared to the remaining samples (Set 2; note that the full Set 2 sample group was used here, although AZGP1-and MFAP5-specific data could also be considered).

In the transcriptomics study, AZGP1 showed higher expression in MLH compared to MUO and MHO samples (Set 1), whereas MFAP5 exhibited lower expression in MLH compared to MHO samples (Set 2).

Notably, the MFAP5 secretion trend observed toward the WAT_NE samples in the proteomics data positively correlates with the MFAP5 transcript expression trend in the transcriptomics study when progressing from MUO to MLH samples – abundance and expression is decreasing when going towards WAT_NE and from MOU to MHL, respectively.

Conversely, AZGP1 displayed increased transcript expression when transitioning from MUO and MHO to MLH samples and a decreasing secretion trend when going towards the WAT_NE samples. These observations suggest that both molecules are subject to context-dependent regulation at multiple biological levels, potentially reflecting differences in transcriptional control, secretion dynamics, or post-translational regulation. This, in turn, raises additional research questions regarding the mechanisms governing their regulation and functional roles under distinct metabolic and physiological conditions. Moreover, these findings serve as a reminder that annotation of a molecule as “secreted” does not necessarily imply active secretion under all biological conditions.

As a final note, when generating sample-rankings and trendlines with clear sample-group separations, as in test case 2, BioTrendFinder will often identify many of the same molecules detected by standard group-based statistical comparisons. However, BioTrendFinder enables users to integrate such statistical signals with expression patterns, annotation, and functional graph information, thereby elevating large sets of identified molecules into ranked lists of candidate targets and effectively reducing the search space for downstream experimental validation. Furthermore, in the examples presented here, we focused on statistically significant trendlines. However, BioTrendFinder is also designed to highlight and explore patterns that fall below formal significance thresholds, enabling researchers to uncover potentially meaningful biological trends that might otherwise be overlooked.

## Method

The BioTrendFinder application has been made in R v. 4.1.3 [59] using Shiny v. 1.7.5 [60] as the main programming language and package, respectively. In addition, BioTrendFinder uses Python v. 3.10.6 for certain tasks described below. BioTrendFinder is deployed at shinyapps.io with the webaddress: https://cphbat.shinyapps.io/biotrendfinder/. In Table 5 the most important R and Python packages can be found.

**Table 5.** R and Python packages used in BioTrendFinder.

| R packages | Python packages |
| --- | --- |
| Plotly [61], shiny [60], shinyjs [62], stringr [63], reticulate [64], DT [65], visNetwork [66], RSQLite [67], DBI [68], data.table [69], future [70], promises [71] | NetworkX [72], Numpy [73], Pandas [74], Matplotlib [75] |

### Upload data

#### 1. Dimensionality reduction

The dimensionality reduction with BioTrendFinder can be performed using a PCA and UMAP. With UMAP, only the first two dimensions are used.

#### 2. Extracting group data from column names

The automatic extraction process yields the structure shown in **Table 6**.

#### 3. Mapping of molecular IDs to STRING IDs

All molecular IDs are mapped to a string ID that captures a wealth of different ID types ranging from gene symbols, Entrez and Ensembl gene IDs to Ensemble transcript and protein IDs as well as UniProt accession numbers and more (**Supplementary table 1**).

### Rank data

#### 4. Sample-ranking visualizations

##### Generating sample-rankings

To automatically generate sample rankings, a half circle consisting of 100 points is constructed around the two-dimensional coordinates of the dimensionality-reduced data, where the circle center is given by the coordinates:

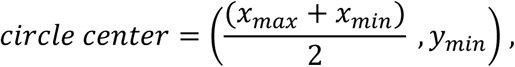

and the circle radius is found by:

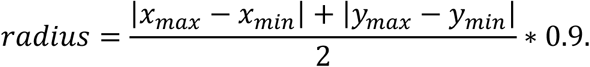

Based on 100 points along the half circle, ranking lines are constructed through these points to derive sample-rankings.

##### Molecule-specific correlation

Each identified sample-ranking is used to compute molecule-specific Spearman coefficients using the cor function from the R stats package.

##### Correlation distribution

The distributions of Spearman correlation coefficients for all molecules are computed across all identified sample-rankings using the `cor` function from the R `stats` package. For each sample-ranking, the Spearman correlation coefficients are grouped into predefined bins spanning the interval from −1 to 1: [−1, −0.8], (−0.8, −0.6], (−0.6, −0.4], (−0.4, −0.2], (−0.2, 0], (0, 0.2], (0.2, 0.4], (0.4, 0.6], (0.6, 0.8], and (0.8, 1].

##### STRING PPI connectivity distribution

For each sample ranking, STRING connectivity data are recorded for the top 100 molecules with the highest and lowest Spearman correlation coefficients (**Table 7**). To evaluate whether the observed number of edges among these molecules exceeds what would be expected by chance, a background edge count is estimated by computing edge counts from 10,000 random draws of 100 nodes. Statistical enrichment of connectivity is expressed as log₂(observed edge count / background edge count) (the ‘Score’ in **Table 7**) .

**Table 6.** Extracting group data. . The words divided by an underscore in the sample names will be combined to form all possible sample-groups. The initial names of the new group columns contain an ‘h’ and a number. These can be changed when clicked.

| Sample_names | h1 | h2 | h3 | h4 | h5 | h6 |
| --- | --- | --- | --- | --- | --- | --- |
| 21_type1_Control | 21 | type1 | Control | 21_type1 | 21_Control | type1_Control |
| 20_type1_Treated | 20 | type1 | Treated | 20_type1 | 20_Treated | type1_Treated |
| 19_type1_Treated | 19 | type1 | Treated | 19_type1 | 19_Treated | type1_Treated |
| 19_type2_Control | 19 | type2 | Control | 19_type2 | 19_Control | type2_Control |
| 21_type2_Treated | 21 | type2 | Treated | 21_type2 | 21_Treated | type2_Treated |
| 20_type2_Control | 20 | type2 | Control | 20_type2 | 20_Control | type2_Control |

**Table 7.** STRING connectivity data.

| **Abbreviations** | **Description** |
| --- | --- |
| Score | The $\log_2$ ratio of the observed edge count to the background edge count. |
| all_edges | Total edge count. |
| neg_nodes | Node count of negatively correlating molecules. |
| neg_edges | Edge count of negatively correlating molecules. |
| neg_singl | Unconnected negatively correlating molecules. |
| pos_nodes | Node count of positively correlating molecules. |
| pos_edges | Edge count of positively correlating molecules. |
| pos_singl | Unconnected positively correlating molecules. |

**Table 8.** Group division scores and categories.

| Score | Percentile | Strength categories |
| --- | --- | --- |
| $[0 - 0.25[$ | $[0\% - 30\%[$ | Negligible |
| $[0.25 - 0.43[$ | $[30\% - 50\%[$ | Weak |
| $[0.43 - 0.63[$ | $[50\% - 70\%[$ | Moderate |
| $[0.63 - 0.85[$ | $[70\% - 90\%[$ | Strong |
| $[0.85 - 1.0[$ | $[90\% - 100\%[$ | Very strong |

***Note***: If the analyzed dataset contains fewer than 100 molecules, STRING PPI connectivity score may be low.

#### 5. Confidence score

The **group confidence score** quantifies how evenly spaced the samples of a given group are along a ranking axis. Let 𝑛 denote the number of samples in the group, and 𝑚 denote the total number of samples. Each sample is assigned a positive integer position along the ranking, 𝑑_obs,j_ ∈ {1,2, …, 𝑚}, reflecting its relative order. By construction, these positions are strictly increasing within the group.

The ideal evenly spaced positions for a group is defined as:

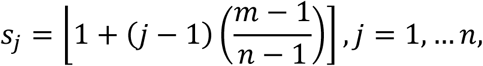

where ⌊⋅] denotes rounding to the nearest integer. The differences between consecutive positions are then computed for both the observed and ideal rankings:

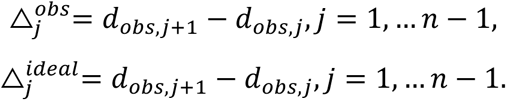

The product of these differences reflects the spread of the group along the ranking:

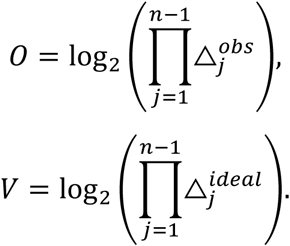

Finally, the group confidence score is defined as

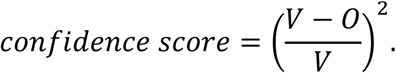

Because the positions are positive integers, the product of the differences is maximized by the evenly spaced configuration, guaranteeing that 0 ≤ O ≤ V and thus 0 ≤ confidence score ≤ 1. A confidence score close to zero indicates that the group’s samples are nearly evenly distributed along the ranking axis, whereas higher values approaching one indicate that the group samples are clustered together, deviating strongly from an even spacing.

### Analyze

#### 6. Spearman coefficient

The Spearman coefficients of the trendlines are computed using the `cor` function from the R `stats` package. Values returned as `NA` are set to zeros.

#### 7. Slope value

BioTrendFinder calculates a trendline score called the **slope value**, which is a modified version of the slope obtained from a linear model fitted using the `lm` function from the R `stats` package. For each molecule, a slope, denoted *a*, is obtained from a linear regression of the molecular trendline. To capture both the magnitude of the trend and the goodness-of-fit of the linear model, the slope value is computed as:

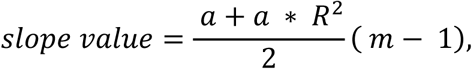

where 𝑅^2^ is the coefficient of determination of the fitted linear model, and 𝑚 is the number of samples.

#### 8. Combined score

The combined score is calculated as the product of the selected score types. Users can choose any combination of the following: Spearman correlation, slope, or mean expression values. For each molecule, the selected scores are multiplied. The sign of the combined score is determined by the first selected score type that contains both positive and negative values.

### Statistics

#### 9. Comparisons of trendline segments and groups

##### Welch test

The p-values of the comparisons are computed using `t.test` function from the R `stats` package with the default settings. Before using the function, a check is made to ensure that none of the variances of the input distributions equals zero. Failing this check will output a value of one.

##### Log2FC

The log2 fold change is calculated by 𝑙𝑜𝑔_2_𝐹𝐶 = log_2_(𝒅) − log_2_(𝒍), where 𝒅 and 𝒍 are the compared distributions in a vector form.

##### Pi-score

The 𝜋-score (𝜋-value) is calculated by 𝜋*-score* = −𝑙𝑜𝑔10(𝑝𝑣𝑎𝑙) × 𝑙𝑜𝑔2𝐹𝐶, where 𝑝𝑣𝑎𝑙 is a given p-values [17].

#### 10. Bundle up

Let 𝑫 ∈ ℝ^n×m^ be the matrix containing the trendline data, 𝑛 is the number of molecular trendlines and 𝑚 is the number of ranked samples. Let 𝑠 be the sign of the mean slope value of the given trendline set.

A column-mean vector 𝒄 = (𝑐_1_, … 𝑐_m_) is computed as:

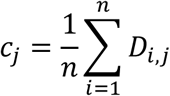

The column means are then adjusted according to:

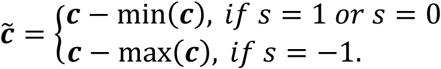

The adjusted matrix 𝑫^-^, which represents the ‘bundle-up’ data, is defined row-wise as:

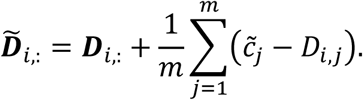

#### 11. Group division scores

Samples along a trendline can fall into one of three categories: left side, right side or excluded from the trendline. To quantify the magnitude of group division trends, we defined a group division score as

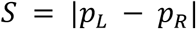

where 𝑝_L_ and 𝑝_R_ represent the proportions of samples assigned to the left and right sides of the trendline, respectively. To establish a reference for interpreting this score, we generated an empirical null distribution using a Dirichlet model. Specifically, we simulated 100000 three-part compositions (𝑝_L_, 𝑝_R_, 𝑝_E_), representing left, right and exclusion proportions, from a Dirichlet distribution with parameters (𝛼_L_, 𝛼_R_, 𝛼_E_) = (1,1,0.2), reflecting symmetry between left and right assignments and a lower expected proportion of exclusions. For each simulated composition, the group division score 𝑆 was computed, and the resulting null distribution was used to define percentile-based strength categories. These categories provide a descriptive measure of deviation from symmetry rather than formal hypothesis testing, allowing us to interpret scores as “Negligible”, “Weak”, “Moderate”, “Strong” or “Very strong” relative to expected variation under a symmetric null model (Table 8).

#### 12. Outlier removal

Outlier removal for an increasing trendline is performed as follows. Let 𝒕 = (𝑡_1_, …, 𝑡_m_) be a trendline, where 𝑚 is the number of ranked samples. Let 𝒂 = (𝑎_1_, … 𝑎_k_) and 𝒃 = (𝑏_l_, …, 𝑏_m_) be two disjoint segments of 𝒕, with 𝑘 < 𝑙 ≤ 𝑚. The segments 𝒂 and 𝒃 are not necessarily contiguous.

Let 𝑧 be a predefined threshold. Let 𝑟_a_ and 𝑟_b_ denote the maximum number of values that may be removed from 𝒂 and 𝒃, respectively. To preserve a minimum segment size, 𝑟_a_ ≤ |𝒂| − 3 and 𝑟_b_ ≤ |𝒃| − 3. The outlier removal procedure iterates over the trendline for at most 𝑟_a_ + 𝑟_b_ iterations.

At each iteration, it is first checked whether:

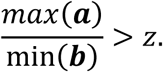

If this condition is false, the iteration stops. If it is true, the following condition is evaluated:

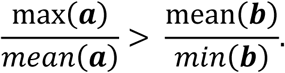

If the condition holds, the value in 𝒂 equal to 𝑚𝑎𝑥(𝒂) is removed from 𝒂. Otherwise, the value in 𝒃 equal to 𝑚𝑖𝑛(𝒃) is removed from 𝒃.

When 𝑧 = 1 (the default value), the procedure removes outliers until no values in the lower segment of the trendline exceed those in the upper segment, or until the maximum number of iterations is reached.

### Functional PPI

#### 13. Louvain community detection

BioTrendFinder utilizes the `louvain_communities` function from the Python package NetworkX to detect Louvain communities. Before community detection is initiated, graph edge weights are transformed such that high values correspond to short distances. Specifically, each weight 𝑤 is converted to 𝑤^r^ → 𝐾 − 𝑤, where 𝐾 is a constant chosen such that 𝐾 > 𝑚𝑎𝑥(𝑤).

#### 14. Ontology edge scores

When enriching graph nodes (molecules) with ontology terms, all applied terms contribute equally; each is assigned an edge weight of 1 / (𝑡𝑜𝑡𝑎𝑙 𝑛𝑢𝑚𝑏𝑒𝑟 𝑜𝑓 𝑖𝑛𝑐𝑙𝑢𝑑𝑒𝑑 𝑡𝑒𝑟𝑚𝑠). If two nodes co-occur in *k* different functional terms, the weight of their connecting edge becomes 𝑘 / (𝑡𝑜𝑡𝑎𝑙 𝑛𝑢𝑚𝑏𝑒𝑟 𝑜𝑓 𝑖𝑛𝑐𝑙𝑢𝑑𝑒𝑑 𝑡𝑒𝑟𝑚𝑠).

#### 15. Combining STRING scores and ontology edge scores

When combining edges from STRING and ontology edges, the edge score from STRING and ontology edge score are combined as well. Because the two scoring systems use different scales, the integrated score is computed as the mean of the **STRING_combined_score** and the ontology edge scores.

#### 16. Set entropy

Set entropy for a given term in the ‘*Advanced selection*’ menu is defined using Shannon entropy to quantify how evenly the term is distributed between two sets. Let

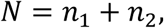

where 𝑛_1_ and 𝑛_2_ are the number of molecules associated with term in Set 1 and Set 2, respectively.

Set entropy is then defined as:

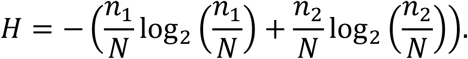

Terms with 𝑛_i_ = 0 are treated as contributing zero to the entropy. Higher entropy values indicate a more even distribution of molecules associated with the given term across the two sets.

### Enrichment analysis

#### 17. Performing enrichment analysis

P-values for each term are computed using the hypergeometric test (`phyper` from the R *stats* package; `lower.tail = FALSE`). Data to perform the test correctly is found in our pre-downloaded STRING database that contains connections between functional terms and STRING IDs. All resulting p-values undergo FDR correction.

#### 18. Preparation of ontology data for graph construction

Knowledge graph data for several of the bio-ontologies used were downloaded from different sources (Table 9) and integrated into the BioTrendFinder back-end. All data was last downloaded 13/11/2025.

**Table 9.**
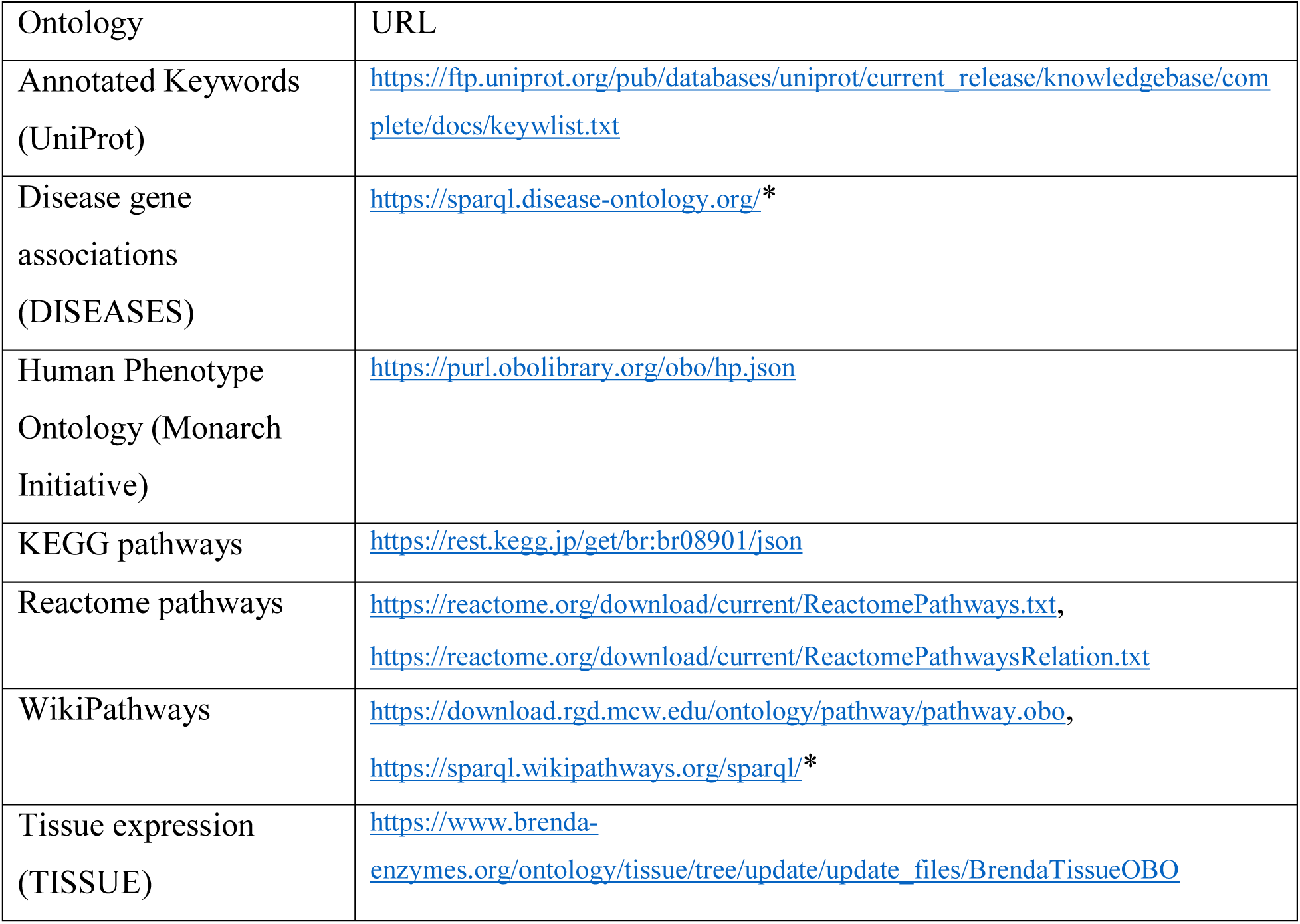
Download links of ontologies. The URLs marked with ‘*’, need additional query data.

All relevant Gene Ontology and Subcellular Localization (COMPARTMENTS) data are retrieved in real time from https://www.ebi.ac.uk/QuickGO/services/ontology/go/terms/graph?, with the allowed relationships limited to ‘is_a’, ‘part_of’, ‘negatively_regulates’, ‘occurs_in’, ‘regulates’, ‘has_part’, and ‘positively_regulates’.

#### 19. Construction of hierarchical graphs

A hierarchical graph for a given bio-ontology is generated using the following procedure. All graphs are visualized using the VisNetwork R library, and all relationships between nodes are defined by the underlying ontology.

1. An undirected graph 𝐺 containing all nodes (i.e., functional terms) is constructed. Edge weights between query nodes (i.e., significant terms) are set to zero, while edge weights between all other nodes are set to one.
2. Based on 𝐺, a subgraph 𝑔 induced by the query nodes is constructed.
3. All connected components of the induced subgraph 𝑔 are identified, and the nodes belonging to each connected component are stored as a set 𝐿 = {𝐶_1_, …, 𝐶_k_}, where 𝐶_i_ denotes the set of nodes comprising the 𝑖𝑡ℎ connected component
4. Shortest paths between each pair of connected components in 𝐿 are computed using Dijkstra’s algorithm (`multi_source_djikstra` from NetworkX).
5. A secondary graph 𝐻 is then constructed in which each node represents a connected component, and edges are weighted by the length of the shortest path between the corresponding components.
6. A minimum spanning tree is computed on 𝐻 to minimize the number of additional nodes required to produce a single connected subgraph 𝑔_final_that includes all query nodes.
7. 𝑔_final_ is converted into a directed graph using the known ontology-defined node relationships.

### Functional module

#### 20. Degree centrality

BioTrendFinder computes a max-normalized degree centrality score, assuming that all nodes belong to the same connected graph. The degree centrality for node 𝑖 is calculated as:

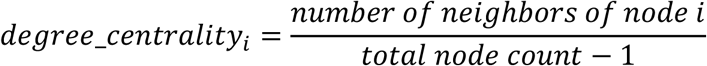

#### 21. Edge weight fraction

For each node, the ontology edge weights associated with that node are summed and divided by the sum of all ontology edge weights in the graph.

#### 22. Node importance

The node importance score for node 𝑖 is calculated as:

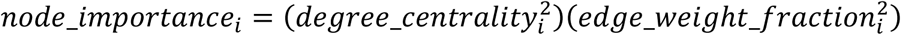

#### 23. Functional module score (FM_score)

The functional module score for a given molecule is defined as the mean of all associated and selected scoring metrics. Prior to averaging, scores are max-normalized by the highest absolute value within each score type. P-values and adjusted *p*-values are first transformed using − log_10_ before max-normalization. A FM_score closer to one indicates higher relevance to the functional module.

### Downloading files and images

#### File download

All tables can be downloaded from the ‘*Download*’ tab as zipped Excel files. BioTrendFinder uses the `rio` R package for file export [76].

#### Downloading graph images

Graph figures in the ‘*Functional PPI*’ and ‘*Enrichment analysis*’ tabs can be downloaded as SVG files. These SVG files are generated using Python (v3.10), NetworkX and matplotlib, based on node coordinates extracted from visNetwork. The underlying graph data can also be downloaded and reused in external graph-editing platforms.

#### Downloading legends separately

Legends for the interactive graphs in the ‘*Functional PPI*’ and ‘*Enrichment analysis*’ tabs are generated in R using ggplot and can be downloaded separately.

#### PlotLy image download

All Plotly-generated figures can be downloaded as SVG files by clicking the download icon in the upper-right corner of the plot. SVG files are vector-based and can be edited using many programs that support PDF or vector graphics formats.

#### Specific figure and table data download

Figure data related to confidence scores in the ‘*Rank data*’ tab and group-division scores underlying figures in the ‘*Statistics*’ tab can be downloaded via buttons located near the respective figures. Additionally, the table in the ‘*Enrichment analysis*’ tab displaying added or missing terms can be downloaded using a button adjacent to the table.

## Author Contributions

AG and CS conceptualized basic aspects of the project. AG further developed and conducted the implementation and analyses. AG and CS wrote and revised the manuscript.

## Data availability

Website is available at: https://cphbat.shinyapps.io/biotrendfinder/. All data and results presented in the manuscript can be downloaded at https://github.com/scheelelab/BioTrendFinder_results_and_data.

## Acknowledgements

Novo Nordisk Foundation Center for Basic Metabolic Research is supported by a donation from the Novo Nordisk Foundation (Grant ID number NNF23SA0084103). This project has received funding from the European Research Council under the European Union’s Horizon 2020 research and innovation programme (grant agreement no. 101002725, to C.S.)

## Supplementary data

**Supplementary table 1:**
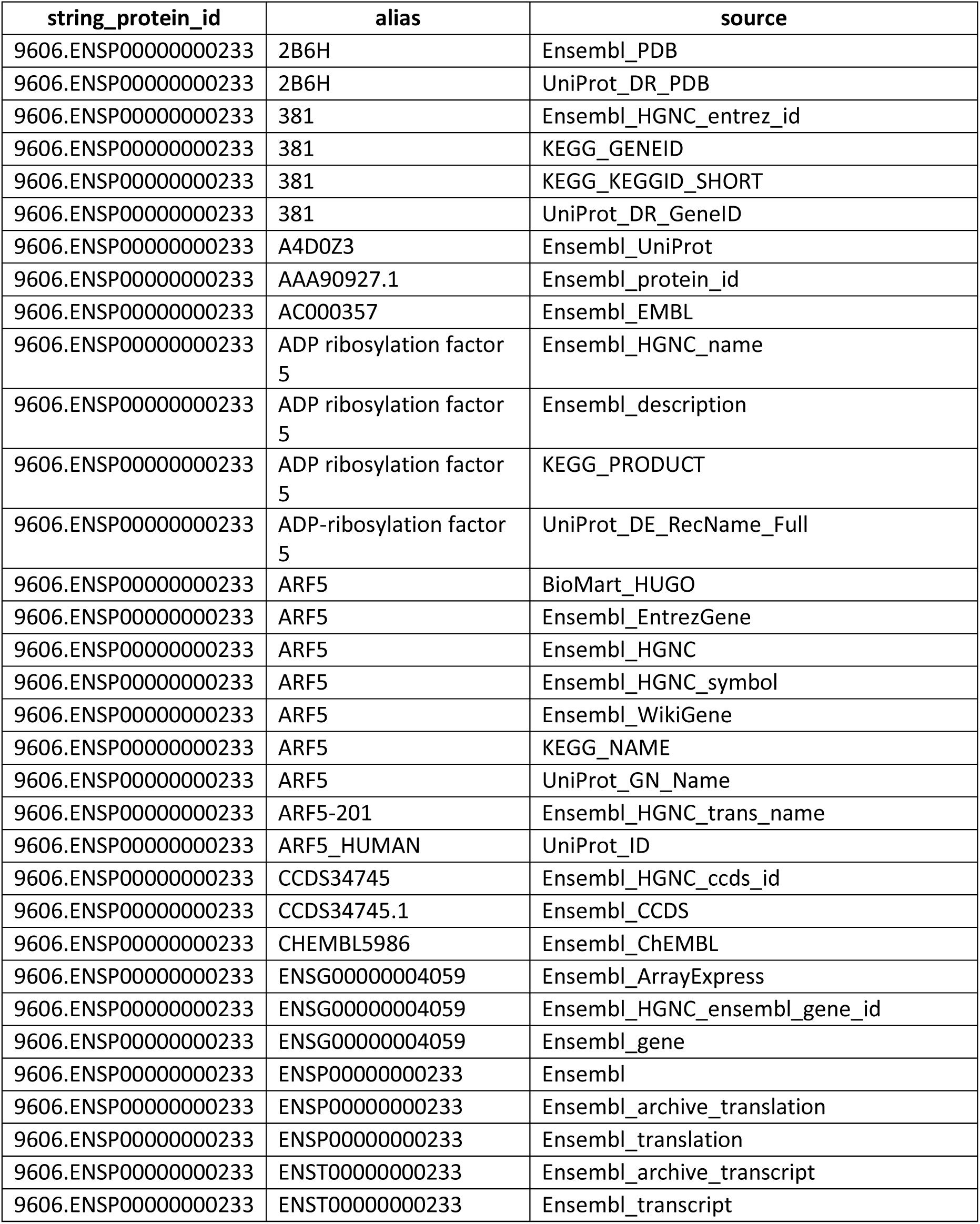
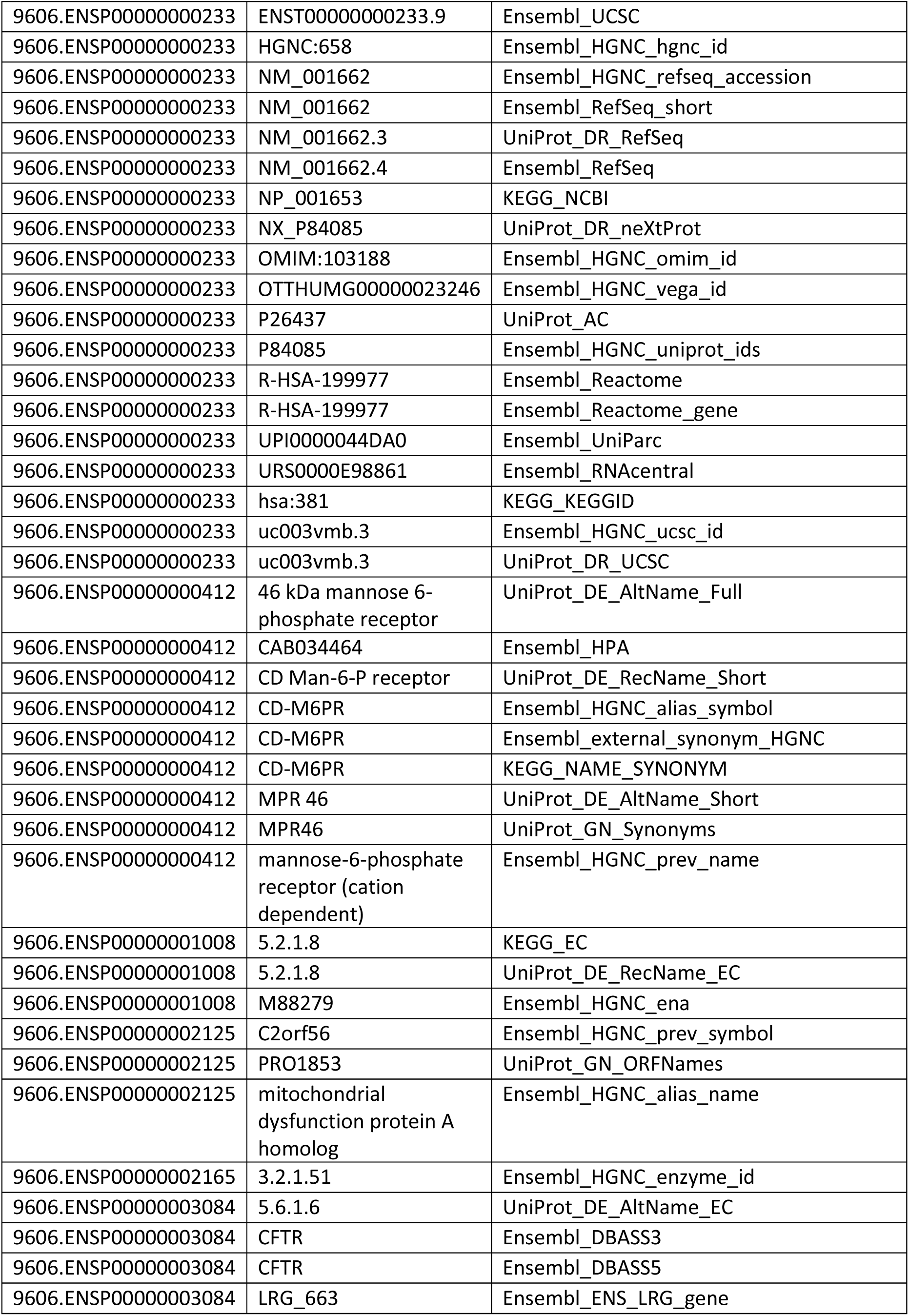
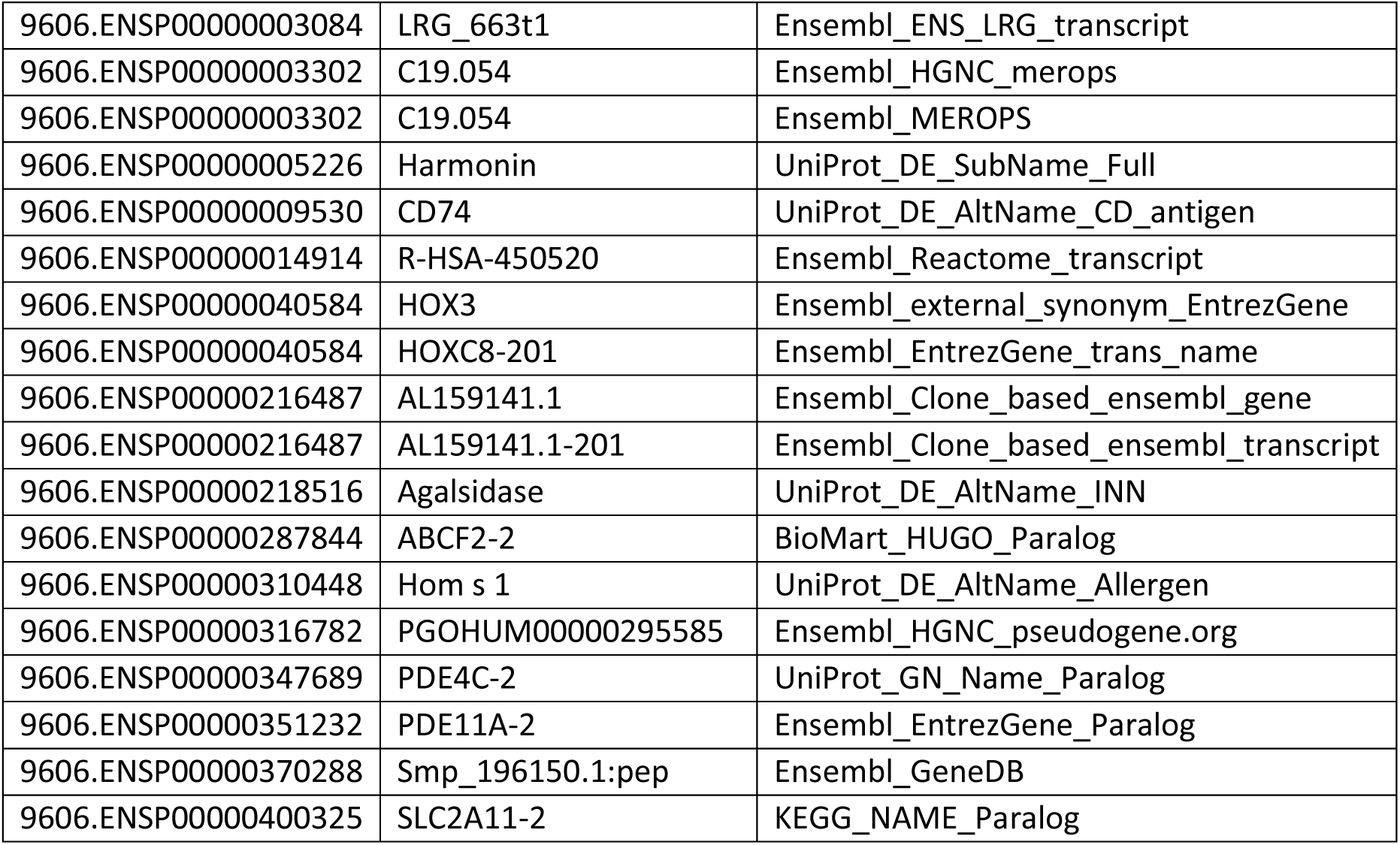
Example of ID conversion table.

## Notes

https://github.com/scheelelab/BioTrendFinder_results_and_data

### Competing Interest Statement

The authors have declared no competing interest.

## References

1. Sainburg T, McInnes L, Gentner TQ (2021) Parametric UMAP Embeddings for Representation and Semisupervised Learning. Neural Comput 33:2881–2907. 10.1162/neco_a_01434

2. Hinzman CP, Singh B, Bansal S, et al (2022) A multi-omics approach identifies pancreatic cancer cell extracellular vesicles as mediators of the unfolded protein response in normal pancreatic epithelial cells. J Extracell Vesicles 11:e12232. 10.1002/jev2.12232

3. Petersen MC, Smith GI, Palacios HH, et al (2024) Cardiometabolic characteristics of people with metabolically healthy and unhealthy obesity. Cell Metab 36:745–761.e5. 10.1016/j.cmet.2024.03.002

4. Anagho-Mattanovich M, Anagho-Mattanovich HA, Argemi-Muntadas L, et al (2025) Multi-omics analysis of thermogenic lipolysis in brown adipocytes. iScience 28:113382. 10.1016/j.isci.2025.113382

5. Szklarczyk D, Kirsch R, Koutrouli M, et al (2023) The STRING database in 2023: protein–protein association networks and functional enrichment analyses for any sequenced genome of interest. Nucleic Acids Res 51:D638–D646. 10.1093/nar/gkac1000

6. Ashburner M, Ball CA, Blake JA, et al (2000) Gene Ontology: tool for the unification of biology. Nat Genet 2000 251 25:25–29. 10.1038/75556

7. Consortium TGO, Aleksander SA, Balhoff JP, et al (2026) The Gene Ontology knowledgebase in 2026. Nucleic Acids Res 54:D1779–D1792. 10.1093/nar/gkaf1292

8. Bateman A, Martin MJ, Orchard S, et al (2025) UniProt: the Universal Protein Knowledgebase in 2025. Nucleic Acids Res 53:D609–D617. 10.1093/nar/gkae1010

9. Grissa D, Junge A, Oprea TI, Jensen LJ (2022) Diseases 2.0: a weekly updated database of disease–gene associations from text mining and data integration. Database 2022:1–8. 10.1093/database/baac019

10. Gargano MA, Matentzoglu N, Coleman B, et al (2024) The Human Phenotype Ontology in 2024: phenotypes around the world. Nucleic Acids Res 52:D1333–D1346. 10.1093/nar/gkad1005

11. Kanehisa M, Furumichi M, Sato Y, et al (2025) KEGG: biological systems database as a model of the real world. Nucleic Acids Res 53:D672–D677. 10.1093/nar/gkae909

12. Milacic M, Beavers D, Conley P, et al (2024) The Reactome Pathway Knowledgebase 2024. Nucleic Acids Res 52:D672–D678. 10.1093/nar/gkad1025

13. Agrawal A, Balcı H, Hanspers K, et al (2024) WikiPathways 2024: next generation pathway database. Nucleic Acids Res 52:D679–D689. 10.1093/nar/gkad960

14. Binder JX, Pletscher-Frankild S, Tsafou K, et al (2014) COMPARTMENTS: unification and visualization of protein subcellular localization evidence. Database 2014:. 10.1093/database/bau012

15. Palasca O, Santos A, Stolte C, et al (2018) TISSUES 2.0: an integrative web resource on mammalian tissue expression. Database 2018:. 10.1093/database/bay003

16. Dyer SC, Austine-Orimoloye O, Azov AG, et al (2025) Ensembl 2025. Nucleic Acids Res 53:D948–D957. 10.1093/nar/gkae1071

17. Xiao Y, Hsiao TH, Suresh U, et al (2014) A novel significance score for gene selection and ranking. Bioinformatics 30:801–807. 10.1093/bioinformatics/btr671

18. von Mering C, Jensen LJ, Snel B, et al (2005) STRING: known and predicted protein–protein associations, integrated and transferred across organisms. Nucleic Acids Res 33:D433–D437. 10.1093/nar/gki005

19. Deshmukh AS, Peijs L, Beaudry JL, et al (2019) Proteomics-Based Comparative Mapping of the Secretomes of Human Brown and White Adipocytes Reveals EPDR1 as a Novel Batokine. Cell Metab 30:963–975.e7. 10.1016/j.cmet.2019.10.001

20. Qiu S, Wu Q, Wang H, et al (2024) AZGP1 in POMC neurons modulates energy homeostasis and metabolism through leptin-mediated STAT3 phosphorylation. Nat Commun 2024 151 15:3377-. 10.1038/s41467-024-47684-9

21. Mracek T, Gao D, Tzanavari T, et al (2010) Downregulation of zinc-α2-glycoprotein in adipose tissue and liver of obese ob/ob mice and by tumour necrosis factor-α in adipocytes. J Endocrinol 204:165–172. 10.1677/JOE-09-0299

22. McClain DA, Sharma NK, Jain S, et al (2018) Adipose Tissue Transferrin and Insulin Resistance. J Clin Endocrinol Metab 103:4197–4208. 10.1210/JC.2018-00770

23. Vaittinen M, Kolehmainen M, Rydén M, et al (2015) MFAP5 is related to obesity-associated adipose tissue and extracellular matrix remodeling and inflammation. Obesity 23:1371–1378. 10.1002/OBY.21103;PAGE:STRING:ARTICLE/CHAPTER

24. Maffei M, Barone I, Scabia G, Santini F (2016) The Multifaceted Haptoglobin in the Context of Adipose Tissue and Metabolism. Endocr Rev 37:403–416. 10.1210/ER.2016-1009

25. Stromsdorfer KL, Yamaguchi S, Yoon MJ, et al (2016) NAMPT-Mediated NAD+ Biosynthesis in Adipocytes Regulates Adipose Tissue Function and Multi-organ Insulin Sensitivity in Mice. Cell Rep 16:1851–1860. 10.1016/J.CELREP.2016.07.027

26. Zhang W, Ren H, Chen W, et al (2025) Nicotinamide phosphoribosyltransferase in NAD+ metabolism: physiological and pathophysiological implications. Cell Death Discov 2025 111 11:371-. 10.1038/s41420-025-02672-w

27. Reynolds TS, Blagg BSJ (2024) Extracellular Heat Shock Protein 90 Alpha (eHsp90α)’s Role in Cancer Progression and the Development of Therapeutic Strategies. Eur J Med Chem 277:116736. 10.1016/j.ejmech.2024.116736

28. Bourboulia D, Woodford MR, Mollapour M (2023) Extracellular HSP90 warms up integrins for an irisin workout. Cell Metab 35:1099–1100. 10.1016/j.cmet.2023.06.002

29. Noormohammadi A, Khodakarami A, Gutierrez-Garcia R, et al (2016) Somatic increase of CCT8 mimics proteostasis of human pluripotent stem cells and extends C. elegans lifespan. Nat Commun 2016 71 7:13649-. 10.1038/ncomms13649

30. Christiansen SF, Haug KBF, Hussain M, et al (2025) Comparative proteomics and micro-RNA analysis of skeletal muscle cell small extracellular vesicles - Unique profiles in cells from severely obese individuals with type 2 diabetes versus normal glucose tolerance. Front Physiol 16:1696916. 10.3389/fphys.2025.1696916

31. Allan MF, Nielsen MK, Pomp D (2000) Gene expression in hypothalamus and brown adipose tissue of mice divergently selected for heat loss. 101152/physiolgenomics200033149 2000:149–156. 10.1152/PHYSIOLGENOMICS.2000.3.3.149

32. Park S, Oh S, Kim N, Kim E (2023) HMBA ameliorates obesity by MYH9- and ACTG1-dependent regulation of hypothalamic neuropeptides . EMBO Mol Med 15:EMMM202318024-. 10.15252/EMMM.202318024/FIGURES/7

33. Guan C, Yang K, Ma C, et al (2025) STING1 targets MYH9 to drive adipogenesis through the AKT/GSK3β/β-catenin pathway. Biochem Biophys Res Commun 749:151352. 10.1016/J.BBRC.2025.151352

34. Faulconnier Y, Boby C, Pires J, et al (2019) Effects of Azgp1−/− on mammary gland, adipose tissue and liver gene expression and milk lipid composition in lactating mice. Gene 692:201–207. 10.1016/J.GENE.2019.01.010

35. Fain JN, Bahouth SW, Madan AK (2004) Haptoglobin release by human adipose tissue in primary culture. J Lipid Res 45:536–542. 10.1194/JLR.M300406-JLR200

36. Gamucci O, Lisi S, Scabia G, et al (2012) Haptoglobin deficiency determines changes in adipocyte size and adipogenesis. Adipocyte 1:142–183. 10.4161/ADIP.20041

37. Zhang Y, Bai X, Shen K, et al (2022) Exosomes Derived from Adipose Mesenchymal Stem Cells Promote Diabetic Chronic Wound Healing through SIRT3/SOD2. Cells 11:2568. 10.3390/CELLS11162568/S1

38. Li Y, Wang T, Li X, et al (2024) SOD2 promotes the immunosuppressive function of mesenchymal stem cells at the expense of adipocyte differentiation. Mol Ther 32:1144–1157. 10.1016/j.ymthe.2024.01.031

39. Golizeh M, Lee K, Ilchenko S, et al (2017) Increased serotransferrin and ceruloplasmin turnover in diet-controlled patients with type 2 diabetes. Free Radic Biol Med 113:461–469. 10.1016/J.FREERADBIOMED.2017.10.373

40. Ma X, Li M, Qiu S, et al (2025) 1MFAP5 enhances the cold resistance of piglets by promoting the transition of adipocyte progenitor cells to fibroblast lineage. J Integr Agric. 10.1016/J.JIA.2025.09.006

41. Zhang T, Li H, Sun S, et al (2023) Microfibrillar-associated protein 5 suppresses adipogenesis by inhibiting essential coactivator of PPARγ. Sci Reports 2023 131 13:5589-. 10.1038/s41598-023-32868-y

42. Yang Loureiro Z, Joyce S, DeSouza T, et al (2023) Wnt signaling preserves progenitor cell multipotency during adipose tissue development. Nat Metab 2023 56 5:1014–1028. 10.1038/s42255-023-00813-y

43. Tang D, Liu Y, Duan R, et al (2024) COL6A6 Peptide Vaccine Alleviates Atherosclerosis through Inducing Immune Response and Regulating Lipid Metabolism in Apoe−/− Mice. Cells 13:1589. 10.3390/cells13181589

44. Drzewicka K, Głuchowska KM, Mlącki M, et al (2025) Chitinase-1 inhibition attenuates metabolic dysregulation and restores homeostasis in MASH animal models. Front Immunol 16:1544973. 10.3389/fimmu.2025.1544973

45. Qian S, Tang Y, Tang QQ (2021) Adipose tissue plasticity and the pleiotropic roles of BMP signaling. J Biol Chem 296:100678. 10.1016/j.jbc.2021.100678

46. Anunciado-Koza RP, Higgins DC, Koza RA (2015) Adipose tissue Mest and Sfrp5 are concomitant with variations of adiposity among inbred mouse strains fed a non-obesogenic diet. Biochimie 124:134. 10.1016/j.biochi.2015.05.007

47. Gill R, Kuo T (2025) Gc inhibition preserves insulin sensitivity and reduces body weight without loss of muscle mass. JCI Insight 10:. 10.1172/jci.insight.195341

48. Akter R, Afrose A, Sharmin S, et al (2022) A comprehensive look into the association of vitamin D levels and vitamin D receptor gene polymorphism with obesity in children. Biomed Pharmacother 153:113285. 10.1016/j.biopha.2022.113285

49. Kolodziejski PA, Pruszynska-Oszmalek E, Micker M, et al (2018) Spexin: A novel regulator of adipogenesis and fat tissue metabolism. Biochim Biophys Acta - Mol Cell Biol Lipids 1863:1228–1236. 10.1016/j.bbalip.2018.08.001

50. Zeng B, Shen Q, Wang B, et al (2024) Spexin ameliorated obesity-related metabolic disorders through promoting white adipose browning mediated by JAK2-STAT3 pathway. Nutr Metab 2024 211 21:22-. 10.1186/s12986-024-00790-3

51. Kreis N-N, Friemel A, Ritter A, et al In-depth analysis of obesity-associated changes in adipose tissue-derived mesenchymal stromal/stem cells and primary cilia function Check for updates. 10.1038/s42003-025-08986-w

52. Saha A, Rahman I, Roy A, et al (2025) Exploring shared genetic factors and prognostic biomarkers in pancreatic cancer and non-alcoholic fatty liver disease: Focus on hsa-miR-29c-3p and COL11A1 axis. Hum Gene 43:201371. 10.1016/j.humgen.2024.201371

53. Leibowitz SF, Akabayashi A, Wang J (1998) Obesity on a High-Fat Diet: Role of Hypothalamic Galanin in Neurons of the Anterior Paraventricular Nucleus Projecting to the Median Eminence. J Neurosci 18:2709. 10.1523/jneurosci.18-07-02709.1998

54. Baek JH, Kim DH, Lee J, et al (2021) Galectin-1 accelerates high-fat diet-induced obesity by activation of peroxisome proliferator-activated receptor gamma (PPARγ) in mice. Cell Death Dis 2021 121 12:66-. 10.1038/s41419-020-03367-z

55. Tans R, van Diepen JA, Bijlsma S, et al (2019) Evaluation of chitotriosidase as a biomarker for adipose tissue inflammation in overweight individuals and type 2 diabetic patients. Int J Obes (Lond) 43:1712–1723. 10.1038/s41366-018-0225-8

56. Oberauer R, Rist W, Lenter MC, et al (2010) EGFL6 is increasingly expressed in human obesity and promotes proliferation of adipose tissue-derived stromal vascular cells. Mol Cell Biochem 343:257–269. 10.1007/s11010-010-0521-7

57. Landgraf K, Kühnapfel A, Schlanstein M, et al (2022) Transcriptome Analyses of Adipose Tissue Samples Identify EGFL6 as a Candidate Gene Involved in Obesity-Related Adipose Tissue Dysfunction in Children. Int J Mol Sci 23:. 10.3390/ijms23084349

58. Plaza-Diáz J, Robles-Sánchez C, Abadiá-Molina F, et al (2017) Adamdec1, Ednrb and Ptgs1/Cox1, inflammation genes upregulated in the intestinal mucosa of obese rats, are downregulated by three probiotic strains. Sci Reports 2017 71 7:1939-. 10.1038/s41598-017-02203-3

59. R Core Team (2022) R: A Language and Environment for Statistical Computing

60. Chang W, Cheng J, Allaire JJ, et al (2023) shiny: Web Application Framework for R

61. Sievert C (2020) Interactive Web-Based Data Visualization with R, plotly, and shiny. Chapman and Hall/CRC

62. Attali D (2021) shinyjs: Easily Improve the User Experience of Your Shiny Apps in Seconds

63. Wickham H (2022) stringr: Simple, Consistent Wrappers for Common String Operations

64. Ushey K, Allaire JJ, Tang Y (2022) reticulate: Interface to “Python”

65. Xie Y, Cheng J, Tan X (2022) DT: A Wrapper of the JavaScript Library “DataTables”

66. Almende B.V. and Contributors, Thieurmel B (2022) visNetwork: Network Visualization using “vis.js” Library

67. Müller K, Wickham H, James DA, Falcon S (2022) RSQLite: SQLite Interface for R

68. R Special Interest Group on Databases (R-SIG-DB), Wickham H, Müller K (2022) DBI: R Database Interface

69. Dowle M, Srinivasan A (2021) data.table: Extension of ‘data.framè

70. Bengtsson H (2021) A Unifying Framework for Parallel and Distributed Processing in R using Futures. R J 13:208–227. 10.32614/RJ-2021-048

71. Cheng J (2021) promises: Abstractions for Promise-Based Asynchronous Programming

72. Hagberg AA, Schult DA, Swart PJ (2008) Exploring Network Structure, Dynamics, and Function using NetworkX. Python Sci Conf. 10.25080/TCWV9851

73. Harris CR, Millman KJ, van der Walt SJ, et al (2020) Array programming with NumPy. Nat 2020 5857825 585:357–362. 10.1038/s41586-020-2649-2

74. McKinney W (2010) {D}ata {S}tructures for {S}tatistical {C}omputing in {P}ython. In: van der Walt S, Millman J (eds) {P}roceedings of the 9th {P}ython in {S}cience {C}onference. pp 56–61

75. Hunter JD (2007) Matplotlib: A 2D Graphics Environment. Comput Sci Eng 9:90–95. 10.1109/MCSE.2007.55

76. Chan C, Leeper TJ, Becker J, Schoch D (2023) rio: A Swiss-army knife for data file I/O

